# The origin, deployment, and evolution of a plant-parasitic nematode effectorome

**DOI:** 10.1101/2024.01.05.574317

**Authors:** Beth Molloy, Dio S. Shin, Jonathan Long, Clement Pellegrin, Beatrice Senatori, Paulo Vieira, Peter Thorpe, Anika Damm, Mariam Ahmad, Kerry Vermeulen, Lida Derevnina, Siyuan Wei, Alexis Sperling, Estefany Reyes Estévez, Samuel Bruty, Victor Hugo Moura de Souza, Olaf Prosper Kranse, Tom Maier, Thomas Baum, Sebastian Eves-van den Akker

**Author notes:** Correspondence: Sebastian Eves-van den Akker. Current address (Clement Pellegrin).

## Abstract

Plant-parasitic nematodes constrain global food security. During parasitism, they secrete effectors into the host plant from two types of pharyngeal gland cells. These effectors elicit profound changes in host biology to suppress immunity and establish a unique feeding organ from which the nematode draws nutrition. Despite the importance of effectors in nematode parasitism, there has been no comprehensive identification and characterisation of the effector repertoire of any plant-parasitic nematode.

To address this, we advance techniques for gland cell isolation and transcriptional analysis to define a stringent annotation of putative effectors for the cyst nematode *Heterodera schachtii* at three key life-stages. We define 659 effector gene loci: 293 “known” high-confidence homologs of plant-parasitic nematode effectors, and 366 “novel” effectors with high gland cell expression. In doing so we define a comprehensive “effectorome” of a plant-parasitic nematode.

Using this effector definition, we provide the first systems-level understanding of the origin, deployment and evolution of a plant-parasitic nematode effectorome. The robust identification of the comprehensive effector repertoire of a plant-parasitic nematode will underpin our understanding of nematode pathology, and hence, inform strategies for crop protection.

## Introduction

Plant-parasitic nematodes (PPNs) are devastating crop parasites which present a considerable threat to global food security. Every major crop can be parasitised by at least one species of nematode, with collective damage estimated at over 80 billion dollars worldwide (Nicol et al. 2011). The most damaging nematode species are those capable of forming intimate, long-term biotrophic relationships with their host plant – namely, the cyst (genera *Heterodera* and *Globodera*) and root-knot nematodes (genus *Meloidogyne*) (J. T. Jones et al. 2013). These species are capable of eliciting profound changes in plant biology to form a unique pseudo-organ: a feeding site from which the nematode draws all its nutrition (M. G. K. Jones 1981). Through this pseudo-organ, the nematode drains the host of essential nutrients and water. This directly affects the fitness of the host and can result in the death of the plant.

Upon nematode infection, the extent of changes to the plant are vast: nematodes have evolved to suppress the plant immune system (Pogorelko et al. 2020; Derevnina et al. 2021), and to manipulate host cell biology, physiology and development to form the feeding site (Molloy, Baum, and Eves-van den Akker 2023). In cyst nematodes, this includes the arrest of the cell cycle in G2 phase, the fragmentation of the vacuole, the proliferation of numerous organelles (including the smooth endoplasmic reticulum, ribosomes, mitochondria, and plastids), and the dissolution of the cell wall leading to the fusion of hundreds of adjacent cells to form a syncytial feeding organ (Golinowski, Grundler, and Sobczak 1996; Grundler, Sobczak, and Golinowski 1998). In this way, a nutrient sink is formed to sustain the female nematode as it matures and reproduces.

To achieve these changes, nematodes – like other parasites and pathogens – deploy molecular tools called effectors. Effectors can be described as parasite or pathogen-secreted molecules that aid the establishment of disease (Hogenhout et al. 2009). Crucially, in PPNs, these effectors are almost exclusively secreted from two types of specialised pharyngeal gland cells: the subventral glands (SvGs) and the dorsal gland (DG) (Hussey and Mims 1990). Expression of secreted proteins in the gland cells is therefore considered synonymous with proteinaceous effector definition in these species. Changes in the content and activity of the gland cells suggests that the SvGs may be predominantly responsible for effector secretion during the early life-stages, whereas the DG is predominantly active during the sedentary life-stages (Bird 1983; Hussey and Mims 1990).

Despite the clear importance of nematode effectors in feeding site establishment, and hence in the success of nematode pathology, we still have an incomplete understanding of the biological functions, or even identities, of individual effectors (Molloy, Baum, and Eves-van den Akker 2023). Predicting genes encoding effectors represents a major bottleneck for the field (Lovelace et al. 2023).

One property that effectors hold in common is that they are secreted by the pathogen into the host. As a result, the presence of a secretion signal, and the absence of transmembrane domains or ER-retention signals have been the dominant criteria for the prediction of proteinaceous effectors in eukaryotic (and many bacterial) plant pathogens (David S. Guttman, Boris A. Vinatzer, Sara F. Sarkar, Max V. Ranall, Gregory Kettler, and Jean T. Greenberg 2002; Sperschneider et al. 2016). However, differentiating the subset of secreted proteins which actually function as effectors from the full set of pathogen secreted proteins is a perennial challenge. For some plant pathogens, characteristic sequences (e.g. RxLR oomycete effectors (Jiang et al. 2008)), structural motifs (e.g. the MAX (*Magnaporthe* Avrs and ToxB like) fold of ascomycete fungal pathogens (de Guillen et al. 2015)), or even codon usage bias (e.g. positive bias for -AA ending codons in unconventionally secreted cytoplasmic effectors in *Magnaporthe oryzae* (Li et al. 2023)) can be used as an additional criterion for effector identification.

In the case of plant-parasitic nematodes, however, we can take advantage of their unique biology to reveal effector identities - specifically, the presence of specialised gland cells for effector production. This understanding led to the discovery of the DOG-box (Dorsal Oesophageal Gland box), a 6 bp non-coding motif enriched in the promoters of dorsal gland effectors in cyst nematodes (Eves-van den Akker and Birch 2016; Eves-van den Akker et al. 2016). This discovery facilitated the prediction of a superset of putative dorsal gland effectors, a number of which were experimentally validated as dorsal gland expressed genes by *in situ* hybridisation. Subsequently, much research to identify nematode effectors has also taken a genomic approach, with a focus on DNA motifs associated with known gland-cell effector loci in a number of plant-parasitic nematode species (Espada et al. 2018; Masonbrink et al. 2019; Vieira et al. 2018). This approach provides a valuable, non-generic (i.e. unlike secretion signals) criterion for identifying effectors, but it is also limited in two regards. Firstly, the use of distinct motifs by distinct species of nematode leads to a lack of generalisability across nematode species. At the same time, the short length of some motifs, such as the DOG-box, leads to a lack of specificity within the genome of a given nematode species.

Advances in targeted transcriptomics, pioneered by (Maier et al. 2013), have allowed the isolation and transcriptomic analysis of plant-parasitic nematode gland cells. Here, we take advantage of these techniques to generate gland cell-specific RNA-seq libraries for the model cyst nematode, *Heterodera schachtii*, at three key life-stages. In combination with the robust reference genome for *H. schachtii* (Siddique et al. 2022) we used these libraries to define a stringent annotation of 659 effector gene loci: 293 “known” high-confidence homologs of plant-parasitic nematode effectors, and 366 “novel” effectors with high gland cell expression. In doing so, we defined a comprehensive effector repertoire, or “effectorome’’ of a plant-parasitic nematode. Using this comprehensive effector definition, we provide the first holistic understanding of the origin, deployment, and evolution of a plant-parasitic nematode effectorome.

Robust identification of the comprehensive effector repertoire of a given plant pathogen is foundational to understanding its pathology, and hence, can inform the development of resistance to crop diseases (Lovelace et al. 2023). Plant pathogen effectors are also important as a means for investigating fundamental host processes, and as promising targets for biotechnological application (Bedell et al. 2012; Frei Dit Frey and Favery 2021). Taken together, results from this work provide an overview of cyst nematode parasitism which can form the basis of further functional studies of the effectorome, and demonstrate the utility of gland-cell transcriptomics as a method for effector discovery.

## Results

### Targeted transcriptomics of *Heterodera schachtii* gland cells

Gland cells were extracted from three life-stages of *H. schachtii* covering the transition to biotrophy: freshly hatched pre-parasitic second-stage juveniles (ppJ2, i.e. before exposure to the host); parasitic J2s (pJ2, i.e. predominantly motile stages extracted from host tissues); and parasitic J3s (pJ3, i.e. sedentary nematodes engaged in biotrophy, Figure 1A and B). For each stage, mRNA from pools of gland cells was sequenced and aligned to the reference genome of *H. schachtii* (Siddique et al. 2022).

**Figure 1.**
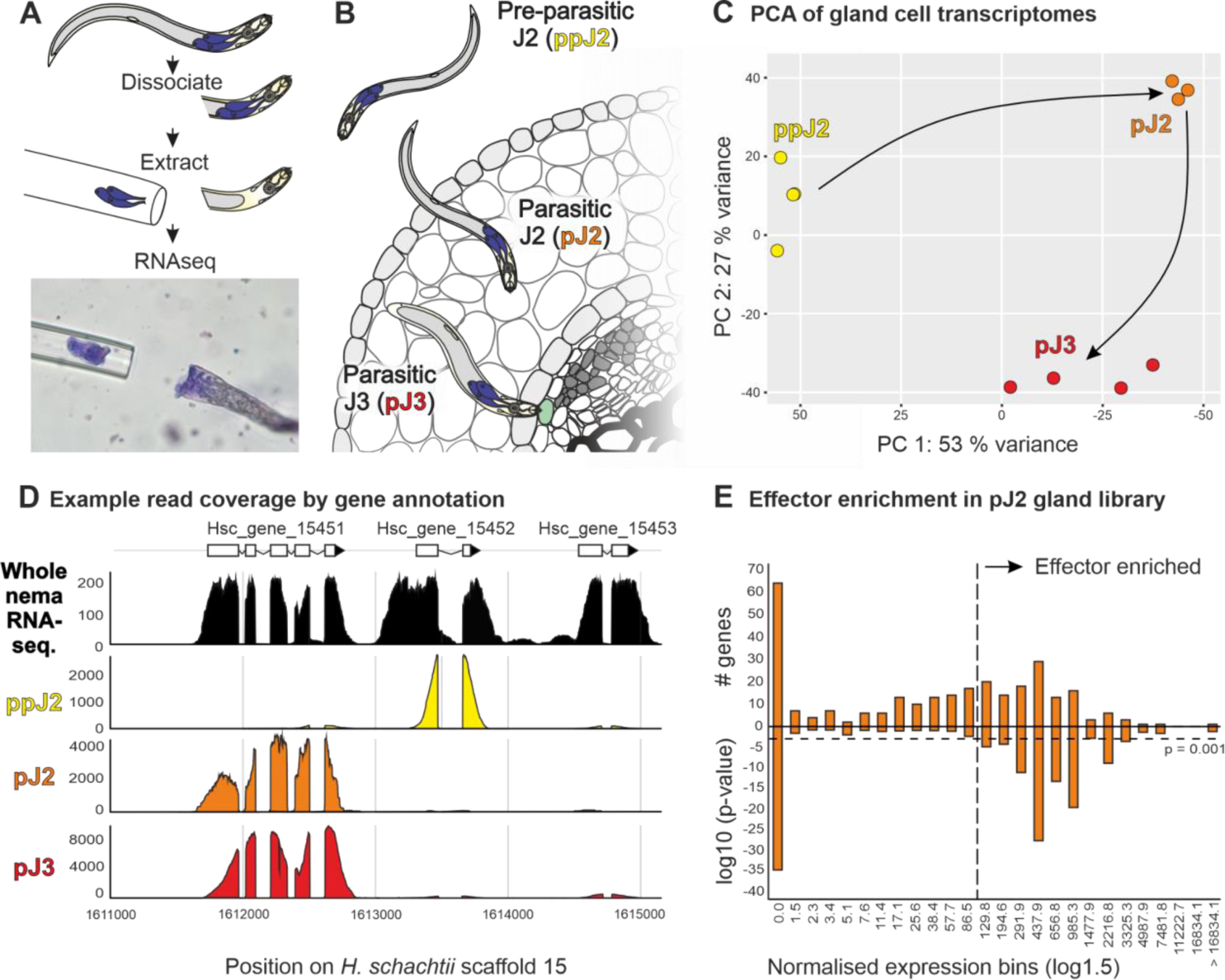
Targeted transcriptomics of *Heterodera schachtii* gland cells. **A)** Schematic representation of gland cell (blue) extraction and sequencing, with representative micrograph below. **B)** Schematic showing the establishment of cyst nematode parasitism, highlighting the 3 life-stages sampled in this study. **C)** Principal components 1 and 2 for gland cell expression gene count data. Circles indicate biological replicates. Arrows indicate progression through the nematode life-stages. **D)** Example coverage of expression data for genes Hsc_gene_15451, Hsc_gene_15452 and Hsc_gene_15453 in the *H. schachtii* genome. The uppermost track shows gene expression data for the whole nematode (black), and the bottom three tracks show expression in gland cell libraries (pre-parasitic J2, ppJ2, yellow; parasitic J2, pJ2, orange; and parasiticJ3, pJ3, red). **E)** Effector enrichment in the parasitic J2 gland cell library (see also Figure S1B). The upper axis shows the number of effector-annotated genes in each expression bin (i.e. at each expression level). Hypergeometric distribution tests were used to determine either the enrichment or depletion of effectors in each bin. The lower axis shows the p-values from these tests. The horizontal dashed line denotes a p-value of 0.001. The vertical dashed line denotes the threshold expression level above which effector genes are consistently enriched.

Gland cell libraries from each life-stage are distinctly different (Figure 1C and 1D), likely capturing changes in expression of the effector repertoire during the transition to biotrophy. As expected, effector-annotated genes and genes encoding putatively secreted proteins (both as defined in (Siddique et al. 2022)) typically have high coverage in these libraries. However, read coverage per gene varies in a continuous distribution. To convert this continuous distribution into a binary classification, all genes were first assigned to one of several expression bins (increasing in a log series) based on how highly they were expressed in a given gland cell library. In general, effector-annotated genes (Figure 1E), and genes encoding putatively secreted proteins (Figure S1A-E), are both statistically depleted in low expression bins and statistically enriched in high expression bins (hypergeometric test, p<0.001). Enrichment of effectors validates the libraries and enrichment of many additional putative secreted proteins suggest that the known effector repertoire is non-exhaustive. Therefore, the lowest expression bin enriched in either - but in most cases both - effector-annotated genes and genes encoding putatively secreted proteins was used to define a cutoff for gland cell expression in each library. Above this cutoff, gland-expressed genes encoding putatively secreted proteins were considered putative effectors (Vieira et al. 2021, 2020; Espada et al. 2018). A majority (64%) of previously effector-annotated genes were recaptured above expression thresholds in gland cell libraries using this approach. Each of the 2,626 gene models above these cutoffs were manually inspected and curated in order to confirm the accuracy of the gene model prior to secreted protein re-prediction to minimise false negatives.

### The *H. schachtii* effectorome

To define a comprehensive effectorome of a plant-parasitic nematode, we combined putative effector identification from the targeted gland cell transcriptomics of ppJ2, pJ2, and pJ3 with effector annotation (largely from (Siddique et al. 2022), but updated with the latest literature). Thousands of gene models were manually examined and curated on Apollo (Lewis et al. 2002) prior to signal peptide prediction to minimise false negatives, maximise their robustness, and ultimately create the highest quality reference database for cyst nematodes with the available data (Table S1). To do this, we retained genes with sequence similarity to previously published effectors if they encode putatively secreted proteins, termed throughout the “knowns”, regardless of their expression in the gland cell libraries (although most are highly expressed), and augmented this with novel putative effectors defined by improbably high gland cell expression, termed the “novels”. Taken together, this combined effector set likely includes most known effectors but underestimates novel effectors (by including only those most highly expressed). Therefore, the combined putative effectorome should be considered a lower bound for a plant-parasitic cyst nematode, comprising 659 loci (293 known, 366 novel) and 774 transcripts (328 known, 446 novel) - Table S1.

The pre-parasitic J2 gland libraries contributed the least, and the parasitic J2 gland libraries contributed the most, to effector prediction (Figure S1F). This is consistent with the expression of known effectors, which typically peak somewhere between 10 and 48 hours post infection (hpi) (Siddique et al. 2022).

The putative effectorome can be grouped into 345 gene “families” based on a combination of common Pfam domains (e.g. the GS-like effectors (Lilley et al. 2018)), pre-computed OrthoMCL (Grynberg et al. 2020) coupled with an all vs all BLAST (for those effectors that do not encode known Pfam domains and cannot be assigned to an orthogroup), or expert knowledge where the nature of the effector precludes the former (e.g. CLEs (Guo et al. 2017) have neither Pfam domains nor function well in BLAST-based analyses due to their short size). The grouping of the putative effectorome into families is highly skewed (Figure 2): the 5 largest families (1.4%) contain a fifth (21%) of all effectors; in contrast, 78% of the families (269/345), and so 41% of all effectors, are the only member of their family in *H. schachtii*. Gland cell specific expression was assigned to putative effectors where it was known for an effector in the same family in a plant-parasitic nematode. Interestingly, the 5 largest gene families are all dorsal gland expressed, and contribute in part to the fact that the DG effectors numerically dominate SvG effectors in the predicted effectorome by nearly 5:1 (Figure 2).

**Figure 2.**
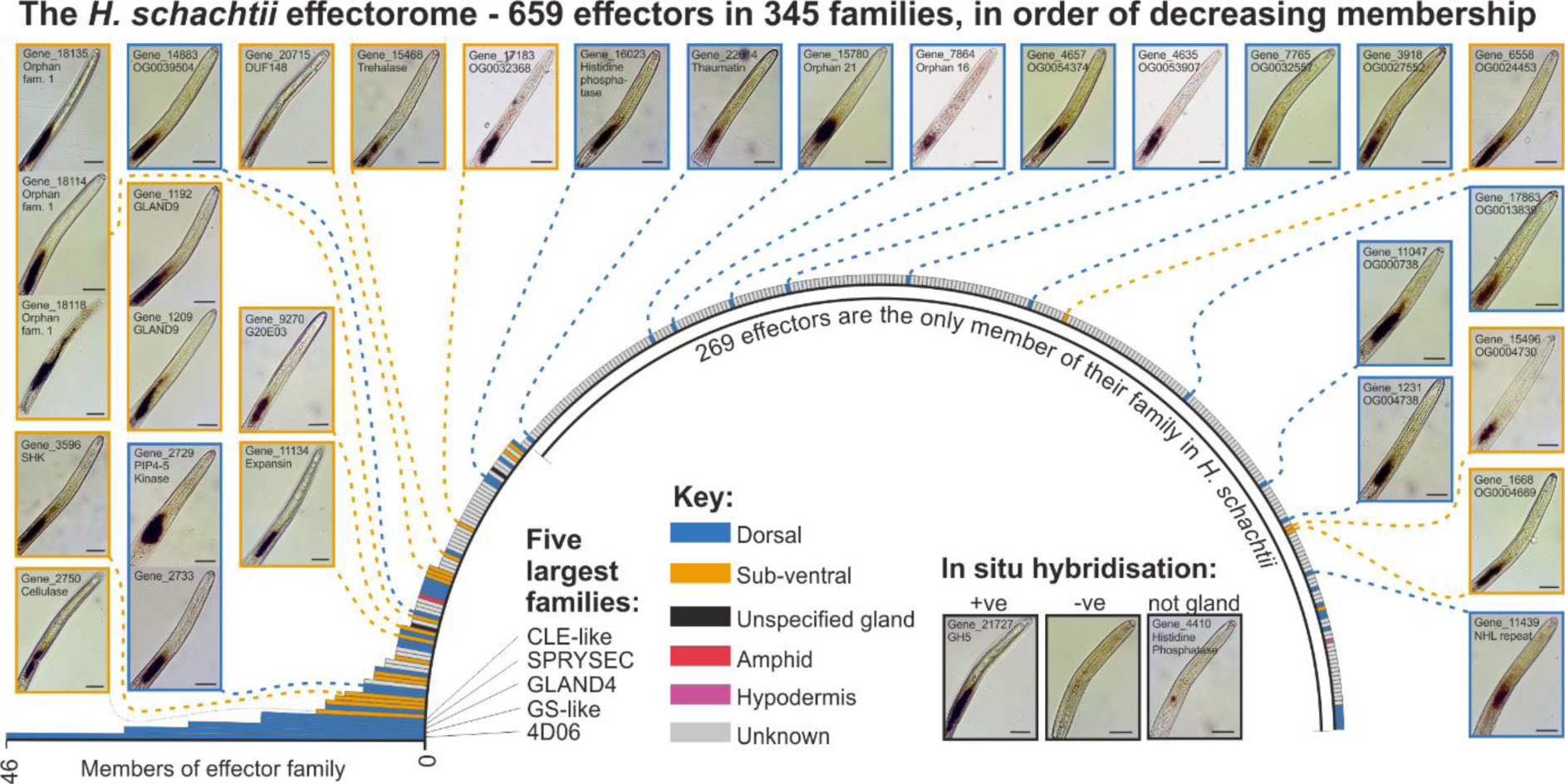
Gene families of the *Heterodera schachtii* effectorome. Radial bar chart showing the number of effectors in each of the 345 families, in order of decreasing membership. The five largest families are named (for additional families see Table S1). Colours indicate gland cell expression of the family, as determined by *in situ* hybridisation, where it is known for at least one member of said family in a plant-parasitic nematode species. Inset panels around the bar chart show *in situ* hybridisations of *H. schachtii* effector transcripts to the gland cells that were produced in this study. Positive (+ve), negative (-ve), and non-gland cell expression (a small unknown cell type adjacent to the glands) are shown. Scale bars represent 25 μm.

We used *in situ* hybridisation, interrogating genes across a range of families, including “knowns” and “novels”, to validate individual genes within the putative effectorome, and to some extent the effectorome as a whole. Of the 31 genes we tested, all but one were confirmed to be gland cell expressed (15 DG, 15 SvG): the exception being Hsc_gene_4410, which was expressed in a small unknown cell type adjacent to the glands, and was subsequently removed from further analyses. Taken together, these data point to the robustness of combined targeted transcriptomics, manual annotation, and the identification pipeline. With this comprehensive and high-confidence putative effectorome in hand, we can now analyse the nature of the effectorome as a whole.

### The effector network

By cross referencing the putative effectorome with the life-stage specific transcriptome (Siddique et al. 2022) we were able to generate a transcriptional network of effectors that elegantly describes the progression of parasitism. Nodes (defined by the 659 putative effector loci), are connected by 9,437 edges (defined by concerted expression across the life cycle above a threshold distance correlation coefficient of 0.975), to reveal a highly connected network (markedly different to a control network of 659 random genes in the genome, Figure S2). Remarkably, all of the connections in the network represent strong positive correlations between effectors. Most effectors have a connection (574/659), and on average are connected to 29 others. Remarkably, this results in one large supercluster, containing 59% of all effectors, connecting those exclusively expressed at pre-parasitic J2 right through to those exclusively expressed during sustained biotrophy (defined as 48 hours post infection to 24 days post infection inclusive (dpi)), by a series of linked subclusters describing the stages in between (Figure 3A). A second separate supercluster contains 23% of effectors, principally those expressed at various individual times between 12 days and 24 dpi. The remaining 18% of effectors are largely independent in the network.

**Figure 3.**
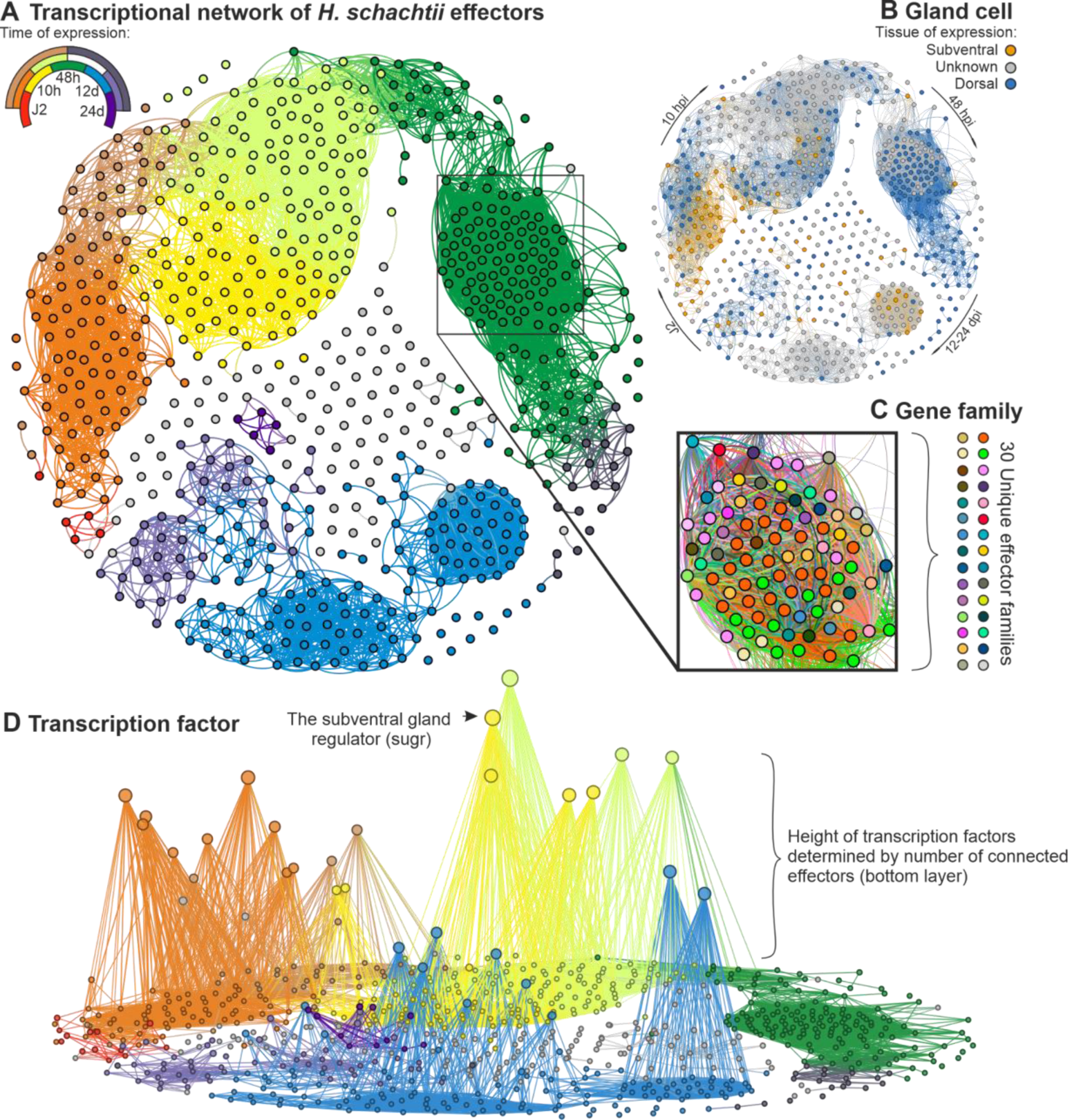
The effector network. **A)** A transcriptional network of *Heterodera schachtii* effectors. Each circle represents one effector locus, and connections between circles indicate a correlation in expression above 0.975 (distance correlation coefficient) across the life cycle. The key indicates the expression supercluster as defined in (Siddique et al. 2022) - where, for example, genes with expression peaking at J2 are shown in red, 10 hours post infection in yellow, and J2_10 hours post infection shown in orange. **B)** The same transcriptional network coloured by gland cell expression (subventral gold, and dorsal blue) of predicted effectors in a given family, where this information is known for at least one effectors in that family in a plant-parasitic nematode **C)** A portion of the 48 hours post infection subcluster coloured by effector family. The key is to illustrate the number of unique families (for specific families therein, see Table S1). **D)** The effector network re-computed with the addition of endogenous nematode transcription factors with at least one connection and at least two non-zero values in gland cell expression libraries. Putative effectors are on the X,Y plane, and transcription factors on the Z axis (height in Z is determined by the number of connections with effectors, with the most connected TFs appearing higher in Z).

We can map various attributes of the effectorome onto the effector network to interrogate both the network itself, and the nature of this parasitism as a whole. For example, we can map expression of effectors in specific gland cell types for effectors in a given family, where this information is known for at least one member of a putative effector family in a plant-parasitic nematode. Mapping gland cell expression to the network in this way supports the accepted view that the subventral gland precedes the dorsal gland during infection: known effectors in the J2 subcluster are exclusively subventral, and known effectors in the 48 hpi subcluster are almost exclusively dorsal (Figure 3B). Given that some times of infection are dominated by a particular gland, this would in principle allow prediction of spatial expression from the network in some (albeit a minority of) cases.

Mapping gene families onto the network reveals the intuitive finding that many of the largest effector families are co-expressed in time. However, in some cases, effector co-expression is more similar between families than within, resulting in individual subclusters that describe a given time being assembled from a diversity of effector families. For example, examining a particularly highly connected part of the 48 hpi subcluster reveals 88 effectors from 30 unique families (Figure 3C).

To understand how effectors from unrelated families are transcriptionally regulated in such a concerted manner, connections between the network and endogenous transcription factors were computed (Figure 3D). Of the 376 transcription factors predicted in the *H. schachtii* genome, 288 show more than 1 non-zero value in the gland cell expression data. Of those, 113 are connected to effectors in the network with a threshold distance correlation coefficient above 0.975 (Figure 3D). Ranking transcription factors in the network by the number of connections they have with putative effectors independently highlights the subventral gland regulator (sugr), a transcription factor that has been shown to regulate the transcription of genes encoding subventral gland effectors, as the second most connected transcription factor (publication to be submitted in parallel). Additional highly connected transcription factors likely regulate other parts of the effector network, and are also highlighted by this approach.

### Evolutionary origins of, and evolutionary pressures on, effectors

To determine when and how effectors evolve, we first cross referenced the putative effectorome with orthologous gene clustering of 61 species, covering the breadth of the nematode phylum and including two outgroup taxa (Grynberg et al. 2020). Effector families were then classified by when the genetic capital (i.e. the underlying genomic sequence) that gave rise to the family is first observed in the phylum. In so doing, two broad classes were identified from the pseudo bimodal distribution (Figure 4A): 1) approximately 20% of effectors (11% of families) are sequence similar to highly conserved genes that predate the nematode phylum (i.e. have a similar sequence in the Tardigrade outgroup), and likely represent duplication and subsequent neofunctionalization (as in the GS-like effectors (Lilley et al. 2018), SPRYSECs (Pearson 2005), peptidases (Robertson, Robertson, and Jones 1999), etc.); and 2) approximately 53% of effectors (61% of families) are only sequence similar to genes that arose since the last common biotrophic ancestor with *Rotylenchulus reniformis*. Very few effectors, approximately 7%, have no similar sequence in any other organism, even in the close sister species *Heterodera glycines*, and are here termed “orphan” effectors.

**Figure 4.**
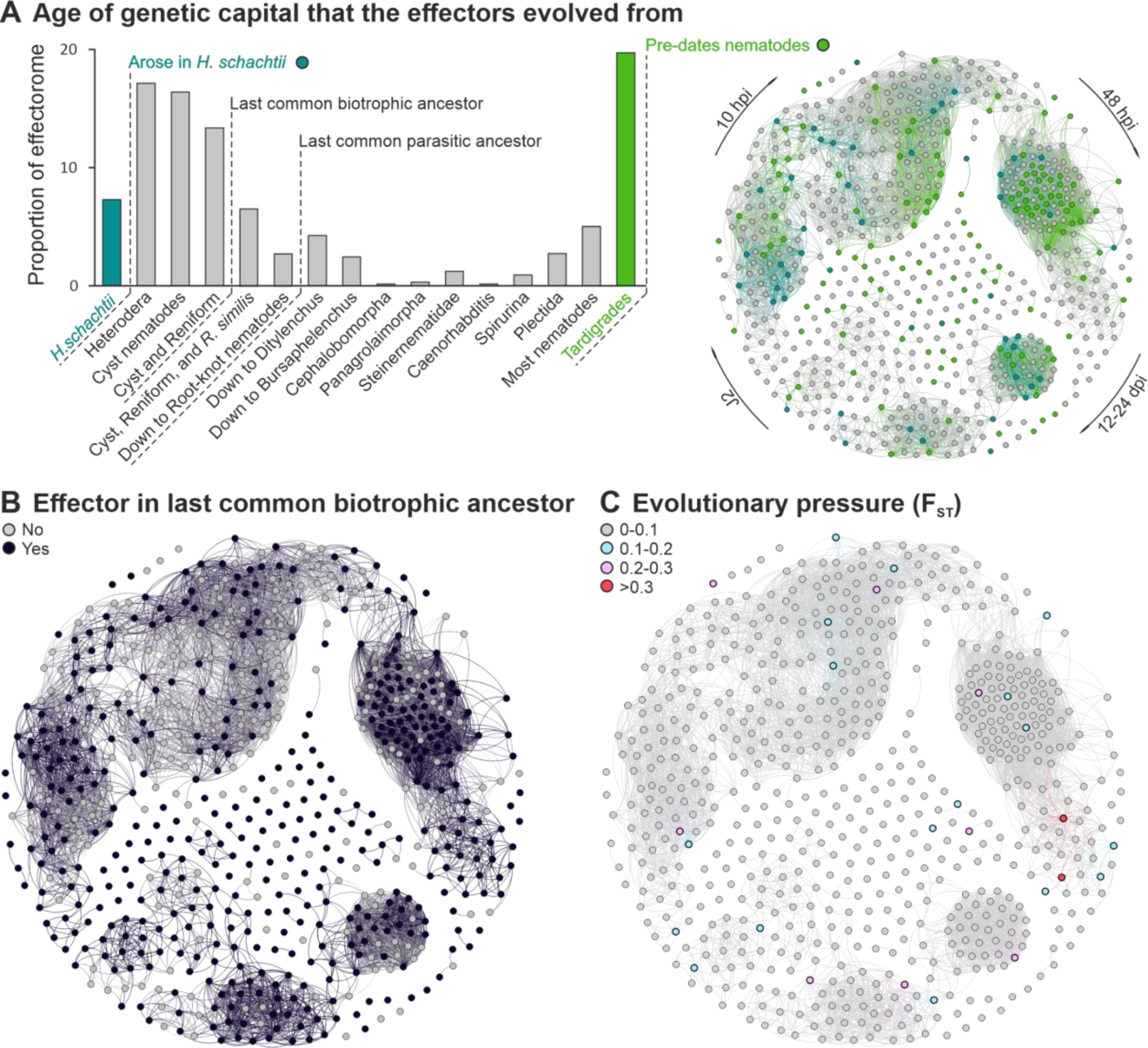
Evolutionary origins of, and evolutionary pressures on, effectors. **A)** A frequency distribution of age of genetic capital (i.e. gene sequence) from which the effectors evolved (i.e. the most distant relative with a similar sequence to a given effector). Two categories are highlighted: those exclusive to *H. schachtii* (teal) and those that pre-date the phylum (green), and mapped to the network (right). **B**) Whether or not each effector is present in an orthogroup with a putative secreted protein from any *Globodera* or *Rotylenchulus* spp. **C**) F_ST_ calculated between *Heterodera schachtii* Bonn and IRS populations, mapped to the network and colour coded by degree.

Contrary to the apparent emerging trend (e.g. 6% of fungal effectors are MAX effectors), we find no evidence of a conserved effector fold, for those effectors that can be predicted at present. We computationally predicted the structures of all proteins in *H. schachtii* and *Globodera rostochiensis* (all models deposited under Dryad accession DOI: 10.5061/dryad.rfj6q57hn). Most effectors are sequence dissimilar to characterised proteins and so result in low-confidence fold prediction. Of the 774 putative effector transcripts identified in this study, 330 transcripts (from 295 loci) fold with an average pLDDT > 50 and pTM > 0.5. A structural similarity network (built using structure-based BLAST, Foldseek (Hutson 2023)) does not identify groups of effectors that would not have otherwise been grouped by sequence similarity and/or expert knowledge of characteristic effector motifs (Figure S4). However, using this approach did reveal unambiguous “hybrid” effectors (Figure S5) resulting from the fusion of two different effector domains, although these hybrid effectors would have been identifiable using sequence similarity alone.

These data demonstrate that the effectorome of *H. schachtii* is assembled from a diversity of genetic capital that itself arose over an extremely long period of time. It does not show when the effector was assembled from said capital. We therefore sought to determine which of the *H. schachtii* effector families were likely already present in the last common ancestor of the cyst nematodes, circa 100 million years ago. We identified orthogroups that contain putative *H. schachtii* effectors, and determined whether the corresponding members of those orthogroups in either *Globodera pallida, G. rostochiensis,* or *R. reniformis* also encode putatively secreted proteins. Using this rough proxy, we estimate that a majority (66%) of effectors (59% of families) were likely present in the last common ancestor of the cyst nematodes (Figure 4B).

Finally, to determine how effectors in *H. schachtii* are currently evolving, we analysed SNPs in sequencing data from two geographically distinct populations: the reference *H. schachtii* “Bonn” population from Germany (Siddique et al. 2022); and the “IRS” population from the Netherlands (van Steenbrugge et al. 2023). We use the fixation index (F_ST_) to determine the genetic differentiation between each gene in the two populations (F_ST_ values range from 0 (no genetic differentiation) to 1 (complete genetic differentiation)). Higher F_ST_ indicates more differentiation between the two populations and hence suggests more evolutionary pressure. As expected, we find that effectors generally have higher F_ST_ values than non-effectors, statistically depleted in the zero F_ST_ bin and enriched in higher F_ST_ bins (hypergeometric test, Figure S3). We mapped F_ST_ to the network as a proxy for positive evolutionary pressure on effector sequences, presumably in large part driven by the host. Mapping F_ST_ to the network shows a relatively even distribution across a diversity of effector families and times of infection (Figure 4C).

### Cross-kingdom gene regulatory network

To determine which genes in the host are transcriptionally co-regulated with nematode effectors, we compared life-stage specific transcriptomic data for the putative nematode effectors and the host plant, *Arabidopsis thaliana* (Siddique et al. 2022). From this we connected each node in the effector network to the host plant genes with correlated expression profiles (defined by concerted expression across infection with a threshold distance correlation coefficient above 0.975). This yielded a network with 6,460 nodes (657 effectors and 5,803 plant genes) and 157,515 edges (20,167 effector-effector connections and 137,348 effector-plant gene connections, Figure 5). The number of connections between effectors and plant genes shows an extremely skewed distribution. The most highly connected effectors, both cellulases, are each connected to 1,011 plant genes (with largely overlapping identities), whilst 50% of effectors are connected to 85 plant genes or fewer. The distribution of connections between plant genes and effectors is similarly skewed, with the most highly connected plant gene connected to 121 effectors, 61.6% of plant genes connected to 10 effectors or fewer, and 18.1% of plant genes connected to just 1 effector.

**Figure 5.**
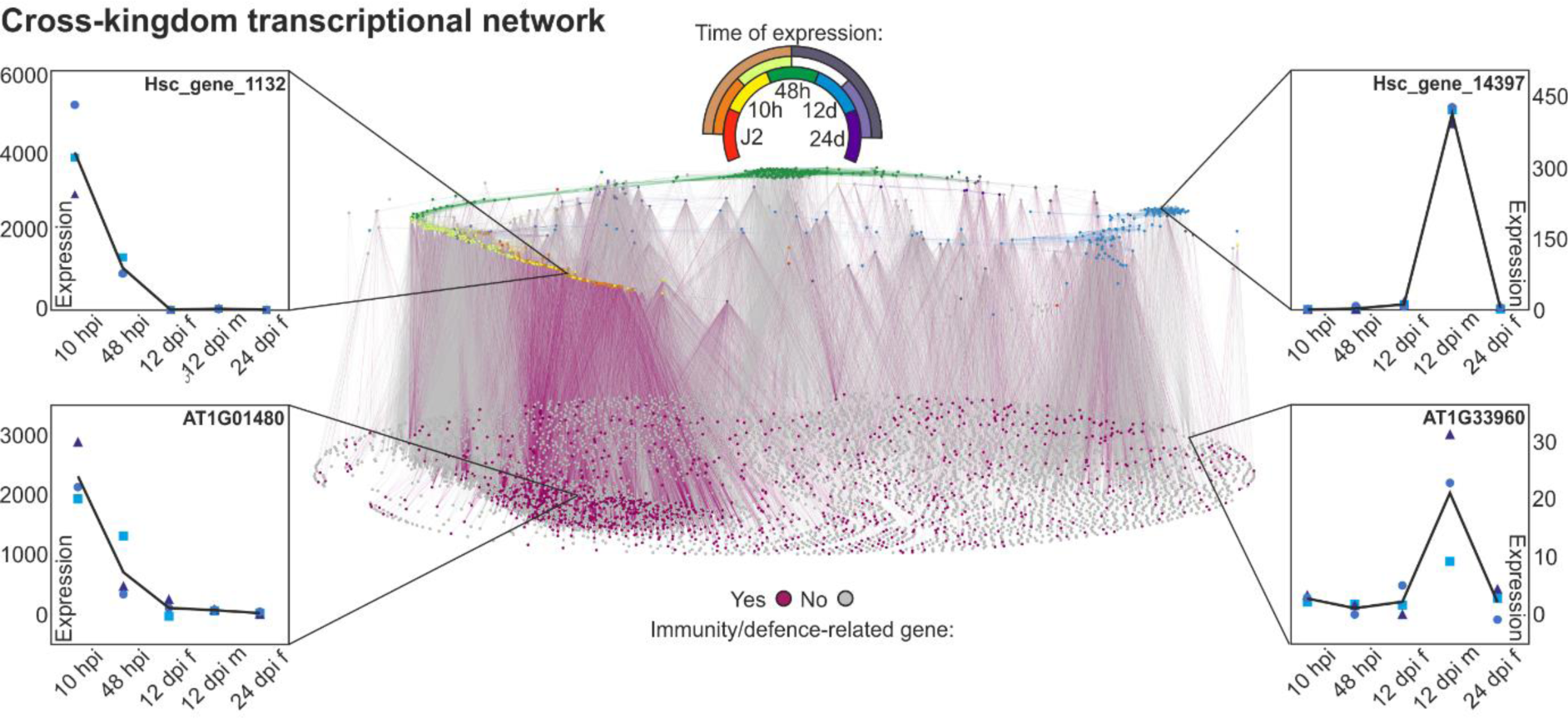
The cross-kingdom transcriptional network. A transcriptional network of *H. schachtii* effector genes and *A. thaliana* genes. In the upper plane, each circle represents one effector locus. In the lower plane, each circle represents a host plant gene. Connections between circles indicate a correlation in expression above 0.975 (distance correlation coefficient) across infection. The upper key indicates the expression supercluster for effectors as defined in (Siddique et al. 2022) and corresponds to the effector plane only. The lower key indicates whether a host plant gene was included in our immune/defence-related gene dataset (purple) or not (grey), and corresponds to the plant genes plane only. Expression profiles (left and right) show gene expression for nematode effectors (top) and plant genes (bottom) with highly correlated expression patterns across the infective life-stages. Individual data points represent biological replicates. Lines represent mean expression across biological replicates.

To understand how effectors might control and/or contribute to host biological processes, for example immunity over the course of infection, we developed a dataset for plant genes present in the cross-kingdom network (i.e. highly correlated in expression with nematode effectors across infection) that are involved in immunity or defence responses. In doing so, we defined an immunity/defence-related gene as: a gene annotated with a GO term in both the GO Slim categories “response to biotic stimulus” (GO:0009607) and “response to stress” (GO:0006950) (according to GO Slim Classification for Plants <u>(Berardini et al. 2004)</u>); or a gene annotated with the GO term “response to wounding” (GO:0009611); or a gene annotated with any other GO term with a name containing the phrases “defence response” or “immune”. In total this dataset contained 1,109 genes, representing 17.9 % of plant genes in the network. To determine when, in the course of the infection, immunity/defence-related genes are co-regulated with effectors, we compared enrichment/depletion of these genes in each of the 29 expression superclusters defined by Siddique et al. (2022). The number of connections between immunity/defence-related genes in the network and effectors in a given supercluster was enriched (1,000 bootstrap, 95 % confidence intervals) mainly in superclusters containing motile stages (i.e. J2 and 10 hpi): “cyst_J2”, “J2”, “10hpi’’, “J2_10hpi”, ‘’48hpi_12dpi_fem.12dpi_male_24dpi’’, and “cyst_J2_10hpi_48hpi”, and depleted in the “cyst”, “48 hpi”, “12dpi_fem.”, ‘’12dpi_fem.12dpi_male’’, “12dpi_fem.12dpi_male_24dpi”, ‘’12dpi_fem._24dpi’’, ‘’increasing’’, and ‘’not clustered but differentially expressed’’ superclusters.

Unlike the effector-effector edges, which represent exclusively positive correlations, 24.6% of effector-plant edges (33,749) are negative correlations. When the “direction” (positive or negative) of effector-plant edges is visualised in the network, positive correlations appear enriched early in infection and evenly distributed at the later life-stages. The vast majority (89.3%) of connections between the effectors and immune/defence-related genes represent positive correlations. These data provide a platform for hypothesis generation, ultimately to accelerate the interrogation of complex plant-nematode interactions.

## Discussion

### Effector identification

Through a combination of targeted transcriptomics and an extensive literature search - both coupled to manual annotation and curation - we have defined a comprehensive putative effectorome of a plant-parasitic nematode. The goal was to retain genes with sequence similarity to previously published effectors, if they encode putative secreted proteins, regardless of their expression in the gland cell libraries (termed the “knowns”), and augment this with novel putative effectors defined by improbably high gland cell expression, (termed the “novels”). While we have been conservative in effector prediction, the number of effectors identified is very large: 659. Using *in situ* hybridisation to test the putative effector prediction resulted in an extremely high true positive rate (30/31 tested effector genes). Coupled with the fact that most (64%), but not all, “known” effectors were above the gland cell transcriptomics threshold, this number is likely a conservative underestimate. If we assume similar numbers of the novels are captured above the threshold, we can estimate this effector repertoire is 80% complete. To put this number in context, more than one in four genes in the secretome, and more than one in forty genes in the genome (2,669 genes, and 26,739 genes respectively (Siddique et al. 2022)) are likely effectors.

Despite progress in identifying promoter motifs associated with gland cell expression (i.e. the DOG Box (Eves-van den Akker et al. 2016)), gland cell targeted transcriptomics appears to be the most efficacious method for effector identification to date. The time of gland cell extraction and, interestingly, the method of analysing the data, each have a very large impact on the number and type of effector identified. The dominant majority of putative effectors identified in this study were identified from the parasitic J2 gland cell library - even if they peak in expression at very different times of infection across the network. The parasitic J3 was the next most informative library, followed by the pre-parasitic J2 which was the least informative (almost uninformative) (Figure S1F).

In all libraries, we used enrichment of effector-annotated genes to identify an expression cutoff above which putatively secreted proteins are likely effectors. Interestingly, using absolute expression values, or relative expression values (gland cell expression divided by whole body expression), does not produce the same results. For the parasitic J2 and J3 libraries, both types of analysis enriched for effector-annotated genes, but captured different sets of putative effectors. For the pre-parasitic J2 libraries only the analysis based on absolute expression values enriched for effectors. When we map those putative effectors identified by each approach onto the network, it is clear that for both the parasitic J2 and the parasitic J3 libraries, enrichment analysis based on relative expression values identified effectors expressed later during infection (Figure S4). We do not fully understand this observation, nor why relative expression-based analyses failed for the pre-parasitic J2, but both may have something to do with the fact that the gland cells themselves increase in size disproportionately to the body post infection, particularly so in the later stages of infection.

Taken together, these data inform future efforts to define effectoromes of plant-parasitic nematodes: gland cell sequencing of parasitic J2 and J3, coupled with absolute and relative expression-based enrichment analyses.

### Deployment

There are two main clusters in the transcriptional network, and at this, albeit arbitrary, threshold they are not connected: one very large supercluster that links effectors expressed at the earliest time point through to those expressed at the latest; and a second large supercluster that principally contains those effectors expressed at various stages 12 dpi onwards. This gap in connectedness most likely reflects the relatively large gap in time between two measurement points (48 hpi and 12 dpi) and not a true biological phenomenon.

Nevertheless, the largest supercluster elegantly highlights the deployment of effectors over time. As previously noted (Siddique et al. 2022) almost no effectors peak in expression at the pre-parasitic J2 stage. There is then a bulge centred on 10 hpi, followed by a second separate bulge at 48 hpi. These data, and indeed the identification of different effectors from each gland cell library (Figure S1F, Figure S4), likely reflect the biology of the system: we speculate that the pre-parasitic J2 nematode cannot express all effectors because it has limited energy reserves; the bulk of the effector repertoire is involved in the very early stages of infection 10-48 hpi, including the transition to biotrophy; once the feeding site is established, a separate, smaller, set of effectors are required for maintenance of the feeding site.

To understand the regulation of this precise control over time, we computed connections to the effector network with transcription factors (TFs) in the parasite genome. In so doing, we inadvertently re-identified the only known regulator of effectors, the subventral gland regulator (sugr, publication to be submitted in parallel). While sugr is the second most connected transcription factor to the effector network, there are many other transcription factors that are highly connected. This will be an exciting area of future study because TFs are also strongly associated with most other times of infection, and this method both identifies, and ranks, these transcription factors by their likely impact on the regulation of the effectorome.

Interestingly, the 48 hpi subcluster has no known transcription factors connected to it. There are several possibilities that may explain this observation. One possibility is that the definition of transcription factor used is imperfect. While this is certainly true, it may or may not be the explanation for this conspicuous absence. Another explanation is that consortia of transcription factors are responsible, each of which does not have a sufficiently similar expression profile to cluster with the group, but together produce the observed pattern. Similarly, it is possible that individual isoforms of a transcription factor would have a highly correlated expression with this group, but that when computed on a per-locus basis do not. All of the above assumes that the expression of the transcription factor/factors itself/themselves do indeed change over time. It is certainly possible that transcription factor(s) that control this group are not regulated at the transcriptional level, but are instead regulated by some other post transcriptional/translational mechanism that gives rise to the observed pattern in the effector transcription without a corresponding pattern of transcription of the factor itself. In any case, understanding the regulation of this subcluster, and indeed the other subclusters, will be the focus of future research.

### Evolution

The effectorome is assembled from a diversity of genetic capital that itself evolved over a very long period of time - approximately 20% of effector sequences are similar to genes that predate the phylum Nematoda. Therefore, caution is advised when analysing effector identification pipelines that exclude genes similar to those in non-parasitic ancestors (i.e. *Caenorhabditis elegans*): if applied to this species they would have missed one in five effectors, including the 2nd and 5th largest families. Generally speaking, dorsal gland effectors tend to be assembled from newer genetic capital, and subventral gland effectors from older genetic capital, although there are many exceptions. The “novel” effectors tend to be assembled from even newer genetic capital (Table S1), which makes sense because many of the known effectors are identified by homology to effectors in another species and so are by definition conserved, at least to some degree. Novels are identified by direct gland cell sequencing and so their identification is not biassed in the same way.

A structural similarity network of predicted effectors did not identify sequence unrelated but structurally similar effectors (Figure S4). This is contrary to the emerging theme (e.g. MAX effectors in ascomycete fungal pathogens (de Guillen et al. 2015)) and suggests that, for *H. schachtii* effectors at least, protein sequence homology captures the dominant majority of relatedness within the effectorome for those that are foldable today with Alpha Fold and ESM fold combined (42%).

Taking a comparative approach, it will be possible to identify a “core” effectorome of the last common biotrophic ancestor of the cyst nematodes, based on these data. While this will require the complete effectorome of at least one other species, carefully selected for its/their position in the phylum, a rough proxy is presented herein based on available genomic data. Here, if an effector had a similar sequence that is also predicted to be secreted in the cyst or reniform nematodes, it was considered to be an effector in said species. Using this information, we can roughly date the emergence of the effector families, and find that many extant families, including the largest, were probably already present in the last common biotrophic ancestor of the cyst nematodes. The conserved ability of the last common biotropic ancestor to manipulate plant development, metabolism, and physiology (as reviewed in (Molloy, Baum, and Eves-van den Akker 2023)) likely resides in extant members of this “core” effectorome.

Intuitively, the older effector families also tend to be the larger effector families. Taken together with the highly skewed membership of effector families, this suggests that the effector repertoire has been moulded by large scale re-shaping/expansion of the “core”, coupled with recent addition of many new small effector families, presumably concurrent with the changes in host/genotype over the same period. Today, evolutionary pressure appears to be remarkably evenly spread across the effector network (both in terms of time of effector deployment but also effector family), there are no obvious discrete or concentrated pressure points. To anthropomorphise, this is probably how the nematode would want it.

### Cross-kingdom regulation

We generated a cross-kingdom transcriptional network for nematode effectors and host plant genes to identify functions that are co-regulated throughout infection. This revealed a highly connected network across all infective stages. By highlighting plant genes of interest in the network (e.g. genes of a particular function or pathway), we can interrogate which plant processes may be altered during parasitism, and when in the life-cycle this might occur.

Importantly, the cross-kingdom network differs from the effector-only network in one key aspect: 25% of the connections are strongly negative (0% of the effector-only network were negative connections). This could possibly reflect suppressive interactions between effectors and plant genes, which is consistent with the fact that nematode effectors have been shown to suppress plant immune responses (Pogorelko et al. 2020; Derevnina et al. 2021). Plant genes with functions in immunity or defence are enriched in the early stages of infection, but the majority of these correlations are positive. This might suggest an initial immune response to the nematode which is later suppressed by nematode effectors. However, we do not know whether the nematodes sampled at 10 hpi were about to successfully infect the plant or not. It is possible that many of these nematodes were unable to overcome the plant immune system, and hence were unable to establish parasitism. While it is tempting to use these data to infer function, this should be tested experimentally before conclusions are drawn.

In contrast, nematodes sampled at the 12 dpi and 24 dpi life-stages are by definition successful, so here we can more confidently hypothesise about roles for effector-plant gene correlations in parasitism. In addition to altering the plant immune system, nematodes also take advantage of plant developmental plasticity to reprogramme many elements of development, physiology and cell biology (Molloy, Baum, and Eves-van den Akker 2023). Where an effector is correlated with genes involved in plant developmental processes in these parasitic clusters, this effector can be a candidate for development altering functions. Uncovering the development altering “toolbox” of plant-parasitic nematodes can enable biotechnology, crop protection, and uncover fundamental aspects of plant biology. This cross-kingdom transcriptional network can provide a basis for identifying potential targets for future functional work in plant-parasitic nematode effector biology.

## Materials and Methods

### Gland cell extraction and sequencing

For the parasitic J2 library generation, gland cells were extracted and library construction proceeded according to previously established methodology for fixed gland cells (Maier et al. 2021). For the pre-parasitic J2 and parasitic J3 library generation, non-fixed gland cells were used in library construction. This was done to improve RNA quality, reduce the number of gland cells needed for input, and reduce the overall rRNA contamination in the final library.

In brief, for each biological replication of each life-stage, nematodes of each life-stage were collected using established methods. 50 μl packed volume of each life-stage were washed in 10 mM MES buffered (pH 6.5) water and resuspended in 100 μl of ice cold 3xHank’s Balanced Salts Solution (14065-056 Gibco-BRL), supplemented with 2% Foetal Bovine Serum (A3160601 Gibco-BRL) and 1 U/μl Superase-in RNAse inhibitor (AM2694 ThermoFisher) (Cutting Buffer). 35 μl of this nematode suspension was transferred into a RNAseZap treated 60 mm glass petri dish and cut with a vibrating razor blade, with the goal of 2-3 cuts per nematode. The cut nematode pieces were recovered by washing the glass dish with 1 ml ice cold Cutting Buffer and transferring this suspension to a 15 ml conical bottom tube on ice. This was repeated until all of the nematode suspension was cut. The contents of the 15 ml conical bottom tube was filtered through a 25 μm tissue filter (Milintyl Biomacs) and into a new 15 ml conical bottom tube, on ice. The cell filtrate was gently pelleted at 1000 g for 3 minutes with a gentle brake. The supernatant above the cell pellet was removed to approximately 100 μl, DAPI was added to a 1:1000 dilution and this was kept on ice. 30 μl of this suspension was transferred onto a coverglass thickness slide, spread across the slide, gently, with a pipette tip and observed and manipulated under an inverted fluorescent microscope with a micromanipulator attached. Using a microinjection needle with a diameter of approximately 20 μm, we microaspirated a total of 10 gland cells (5 dorsal and 5 subventral pairs) into the needle (use fluorescence and DAPI filter to aid in observing gland cells, if needed) utilising CellTram Oil to generate a vacuum. After the completion of the collection of each biological replication, the set of collected gland cells were transferred into a 5 μl drop of IDTE (10 mM Tris, 0.1 mM EDTA) buffer that has been placed in the neck of a 200 μl thin wall PCR tube laid on its side and placed on a fresh coverglass thickness slide. The micromanipulator and CellTram Oil were used to generate back pressure to expel the gland cells from the needle and into the 5 μl drop. This drop was then spun to the bottom of the tube via a tabletop microcentrifuge, flash frozen and placed at −80 °C.

Once all biological replications of glands from each life-stage were collected, the 5 μl samples were used as input into the SMART-Seq v4 Ultra Low Input RNA Kit (Takara Bio USA) (for parasitic J3 library generation) or the SMART-Seq mRNA LP Kit (Takara Bio USA) (for pre-parasitic J2 library generation). We followed the protocol for starting with RNA or Cells Sorted into Non-CSS Buffer. Additionally, we modified the overall protocol to include a cell lysis optimization step prior to First Strand Synthesis, where we performed 3 rounds of freeze-thaw on all samples to improve cell lysis efficiency. Libraries were sequenced using 150 bp paired end reads.

All RNAseq reads were analysed with FastQC v.0.11.8 (“Babraham Bioinformatics - FastQC A Quality Control Tool for High Throughput Sequence Data” n.d.) and trimmed using BBduk v38.34 (*Bbduk.Sh at Master · BioInfoTools/BBMap* n.d.). Only reads with a minimum Phred Quality Score of 20, minimum length of 75 bp, and without adapters were retained. Low quality bases were also removed from the 5’ ends of reads in accordance with FastQC per base sequence quality analysis. Trimmed reads from each library were mapped to the *H. schachtii* 1.2 reference genome (Siddique et al. 2022) using STAR v2.7.10b (Dobin et al. 2013). Mapped reads were visualised using Apollo (Lewis et al. 2002). The htseq-count function of HTseq v0.12.4 (Anders, Pyl, and Huber 2015) was used to count read coverage per gene. For the ppJ2 and pJ3 libraries only, uniquely mapped spliced reads were counted to remove artefacts attributed to the low input library prep method for these two gland cell types. Count tables were loaded into R v4.2.1 using the tidyverse package (Wickham et al. 2019) and normalised using DESeq2 v1.22.1 (Love, Huber, and Anders 2014). The clustering of gene counts from processed RNA-seq data from each biological replicate was visualised by a Principal Component Analysis (PCA) using the plotPCA function in R.

### Effector identification

For each of the three life-stages (ppJ2, pJ2 and pJ3), mean normalised gland cell expression was calculated for each gene. Mean gland cell counts were expressed both as is (termed “absolute expression values’’), and relative to the rest of the body (termed “relative expression values’’) by dividing gland cell expression by whole nematode expression of the corresponding life-stage (e.g. gland cell values for pJ2 were divided by whole nematode values at 10hpi).

Gland cell expression values (absolute and relative) were sorted into ‘bins’ (each bin represents a range of expression values). For each expression bin the enrichment or depletion of 248 predetermined high-confidence *H. schachtii* effector genes from (Siddique et al. 2022) was determined by a hypergeometric test (Figure S1). The minimum expression level at which effector genes were enriched established the threshold expression level above which genes were considered to be putative effectors provided they: i) encode predicted secretion signals (SignalP v4.1); ii) contain no TM domains (TMHMM) or iii) ER retention motifs (Regular expression). The size of expression bins was chosen individually for each life-stage (and for absolute and relative values) based on the ‘hit-rate’ (i.e. the ratio of known effector genes to the total number of genes captured above the threshold) with lower ratios being preferred. For ppJ2 relative values, effectors were not enriched at any expression level for any bin size, and so only absolute values were used for further analysis. Gene models for all 2,626 genes above the respective thresholds for each life-stage (for both absolute and relative where possible) were manually inspected and re-annotated on Apollo (Lewis et al. 2002) where the gene prediction had failed to correctly capture gene structure. For corrected genes, secretion signals and transmembrane domains were repredicted using SignalP 4.1 and SignalP 6.0 (Teufel et al. 2022). Predetermined known effector genes (Siddique et al. 2022) and novel highly gland cell expressed predicted effector genes were combined to form a comprehensive list of predicted *H. schachtii* effector genes (Table S1).

### *In situ* hybridisation

*In situ* hybridizations were performed using ppJ2 of *H. schachtii* following previously published methodology (de Boer et al. 1998). Specific primers were designed to amplify a product for each of the candidate effector genes using a cDNA library produced from ppJ2s (Table S2). The resulting PCR products were then used as a template for generation of sense and antisense DIG-labelled probes using a DIG-nucleotide labelling kit (Roche, Indianapolis, IN, USA). Hybridised probes within the nematode tissues were detected using an anti-DIG antibody conjugated to alkaline phosphatase and its substrate. Nematode segments were observed using a DP73 digital Olympus camera mounted on a Bx51 Olympus microscope.

### Effector family prediction

Orthogroups were assigned to predicted effector genes based on a pre-computed OrthoMCL analysis including 59 species across the phylum *Nematoda*, and 2 outgroup Tardigrade species (Grynberg et al. 2020). Previously, 248 high-confidence effectors were assigned to effector families based on sequence similarity to known plant-parasitic nematode effectors, and the presence of known effector motifs (Siddique et al. 2022). Novel predicted effectors which clustered into the same orthogroups, or shared key functional annotations (Pfam domains) with a known effector (e.g. glutathione synthetase (GS)-like domains (Lilley et al. 2018)) were considered to be in the same effector family. For putative effectors which did not contain characteristic effector Pfam dominas, or share an orthogroup with a known effector, orthogroups were used to define predicted families. Genes with no informative Pfam domains and no assigned orthogroup were compared by sequence similarity (BLAST) to all effector genes. All BLAST alignments were manually inspected and families were assigned accordingly. Expert knowledge was used to assign predicted effector genes to families where the nature of the effector precludes identification by Pfam domains or sequence similarity (e.g. CLEs (Guo et al. 2017) do not have Pfam domains and their short size means they do not function well in BLAST-based analyses). After these combined analyses, genes with no assigned family were considered to be *H. schachtii* specific ‘orphans’. Orphans with sequence similarity to other orphans were assigned to ‘orphan families’.

### Evolutionary origins and pressure

Evolutionary origins of predicted effectors were assigned based on pre-computed OrthoMCL data (Grynberg et al. 2020). Sequences with orthologs in other nematode (or tardigrade outgroup) species were considered to have been present in the last common ancestor shared between that species and *H. schachtii*. Assigned evolutionary origins were then manually curated and updated where expert knowledge contradicted OrthoMCL data or data was absent (e.g. where a GS domain is present, ‘predates nematodes’ was assigned as the sequence origin because the sequence that gave rise to GS effectors predates the phylum *Nematoda* (Lilley et al. 2018)). Orthogroups containing putative *H. schachtii* effectors were identified and corresponding members of those orthogroups from either *G. pallida* (Cotton et al. 2014), Sonawala et al., 2023), *G. rostochiensis* (Eves-van den Akker et al. 2016), or *R. reniformis* (Eves-Van Den Akker et al. 2016) were analysed for the presence of secreted proteins. Secreted proteins were predicted using Signal P 4.1 and the absence of transmembrane domains (using TMHMM). The presence of a secreted ortholog in one of these species was used as a rough proxy for the presence of a homologous effector in the last common ancestor of cyst nematodes.

Genomic Illumina reads from the “IRS” (van Steenbrugge et al. 2023) and “Bonn’’ populations (Siddique et al., 2022) were trimmed of adapters and low quality bases using trimmomatic (HEADCROP:9 ILLUMINACLIP:TruSeq3-PE.fa:2:30:10 LEADING:25 TRAILING:25 SLIDINGWINDOW:10:25 MINLEN:100, (Bolger, Lohse, and Usadel 2014)) and mapped to the reference *H. schachtii* genome using bwa-mem. Duplicates were marked and removed, reads were sorted and read groups added using Picard Tools (version 3.1.0) (http://broadinstitute.github.io/picard/). The deduplicated bam files were then converted to an mpileup file with Q20 threshold using samtools (1.16.1). Popoolation2 (version 1201) was used to convert the file to a sync format and estimate FST values. Scripts and environments can be found here: https://github.com/peterthorpe5/H.schachtii_FST.

### Transcriptional network analyses

Expression profiles of predicted effectors across the nematode life cycle (Siddique et al. 2022) were loaded into R v4.2.1 using the tidyverse package (Wickham et al. 2019)), and pairwise distance correlation coefficients were computed using the energy package (https://CRAN.R-project.org/package=energy). A network of distance correlation coefficients was generated in R v4.2.1 at an arbitrary edge threshold of 0.975. Directionality of correlation was estimated using Pearson’s correlation coefficient. Various attributes were assigned to nodes in the network (expression supercluster (Siddique et al. 2022), gland cell expression (Table S1, assembled from the literature and *in situ* hybridisations in this paper), evolutionary origin and evolutionary pressure (described above), using custom R scripts (https://github.com/BethMolloy/Effectorome_H_schachtii). The *H.schachtii* TFome prediction was based on the Pfam domains found in the *C. elegans* TFome as defined in Kummerfeld&Teichmann, 2006 (DBD database, (Kummerfeld and Teichmann 2006)) and Hu et al, 2019 (AnimalTFDB v3.0, (Hu et al. 2019)) with the addition of PF00105 (Zinc finger, C4 type (two domains)). Predicted transcription factors (TFs) with expression in at least two gland cell libraries were added to the effector network if they shared at least one connection with an effector (above a threshold distance correlation coefficient of 0.975). The number of connections with predicted effectors for each TF was added as a node attribute and used to determine height in the Z axis (https://github.com/BethMolloy/Effectorome_H_schachtii). All networks were visualised using Gephi v0.10.1 (Bastian, Heymann, and Jacomy n.d.).

For the cross-kingdom transcriptional network, we subtracted mean uninfected sample expression values from infected sample expression values at each timepoint across the life-stage specific transcriptome to isolate infection-specific changes in gene expression (Siddique et al. 2022). Distance correlation coefficients were computed between effector genes and normalised plant gene expression profiles. Plant genes were included in the network if they were successfully assigned to a supercluster by (Siddique et al. 2022) and shared at least one connection with an effector (above a threshold distance correlation coefficient of 0.975). Likewise, effector-effector connections were also included in the network, while effector genes with no connections to any other effectors or plant genes were excluded to aid visualisation.

*Arabidopsis thaliana* immunity/defence-related genes were defined as: genes annotated with a GO term in both the GO Slim categories “response to biotic stimulus” (GO:0009607) and “response to stress” (GO:0006950) (according to GO Slim Classification for Plants (Berardini et al. 2004)); or genes annotated with the GO term “response to wounding” (GO:0009611); or genes annotated with any GO term containing the phrases “defence response” or “immune”. A manual inspection of 200 randomly chosen genes from this dataset identified one false positive. Presence or absence of each plant in this immunity/defence dataset was then added as an attribute to the network.

Enrichment or depletion of connections between immunity/defence-related genes and effectors in a given supercluster (as defined in (Siddique et al. 2022) was calculated using a bootstrapping approach. The total number of connections to immunity/defence-related plant genes was counted for each effector supercluster and compared to 1,000 simulations in which immunity/defence-related gene identity was assigned at random to plant genes in the network using R function Sample without replacement (https://github.com/BethMolloy/Effectorome_H_schachtii). Where the true number of immunity/defence-related genes connected to a given supercluster was greater or less than 95 % confidence intervals expected for random assignment of immunity/defence-related genes, immunity/defence-related genes were considered enriched or depleted.

### Protein structure prediction and clustering

Signal peptides were cleaved from amino acid sequences for all secreted proteins in the *H. schachtii* (Siddique et al. 2022), and *G. rostochiensis* genomes (Eves-van den Akker et al. 2016) using SignalP - 4.1 (Petersen et al. 2011)). Sequences were aligned to the ColabFold v1.5.2 (Mirdita et al. 2022; Jumper et al. 2021) database using build in MMseqs2 (Steinegger and Söding 2017). Where effectors were manually annotated and corrected, the corrected sequences were used in place of the original gene predictions. Protein structure was predicted using ColabFold v1.5.2 (Jumper et al. 2021; Mirdita et al. 2022), which has AlphaFold v2.3.1 integrated, with three recycles per model. Predicted folds from the genome with an average pLDDT score < 50 and a pTM score of < 0.5, were discarded. For predicted effector structures, folds below these values were predicted again using ESMfold v1.0.3 (Lin et al. 2023), and filtered and discarded if folds were below the same cutoff quality metric. Structural similarity between effectors was predicted by an all-vs-all search using Foldseek (van Kempen et al. 2023; Barrio-Hernandez et al. 2023) and connections in the similarity network were permitted at TM-scores of >0.5 and above. Relevant scripts are available at: https://github.com/BethMolloy/Effectorome_H_schachtii/tree/main/ProteinFolding.

## Data availability

Raw reads are deposited in ENA accession PRJEB71499 Predicted structures are deposited in DRYAD accession DOI: 10.5061/dryad.rfj6q57hn Transcriptional network files are deposited in DRYAD accession DOI: 10.5061/dryad.rfj6q57hn The predicted Effectorome is available in Table S1.

Scripts unique to this manuscript are deposited under github accessions: https://github.com/BethMolloy/Effectorome_H_schachtii

https://github.com/peterthorpe5/H.schachtii_FST

## Author contributions

Effector annotation - Mariam Ahmad, Beth Molloy,

Data analysis - Beth Molloy, Dio S. Shin, Jonathan Long, Clement Pellegrin, Peter Thorpe, Sebastian Eves-van den Akker

Alphafold - Olaf Prosper Kranse, Samuel Bruty, Anika Damm, Lida Derevnina, Estefany, Reyes Estévez, Victor Hugo Moura de Souza, Beth Molloy, Alexis Sperling, Kerry Vermeulen, Siyuan Wei.

ESMfold, protein structure similarity analyses - Olaf Prosper Kranse

Gland cell sequencing - Beatrice Senatori, Tom Maier and Thomas Baum

*In situ* hybridisation - Paulo Vieira

Writing - Beth Molloy and Sebastian Eves-van den Akker.

## Declarations

### Ethics approval and consent to participate

Not applicable.

### Consent for publication

Not applicable.

---

**Figure S1.**
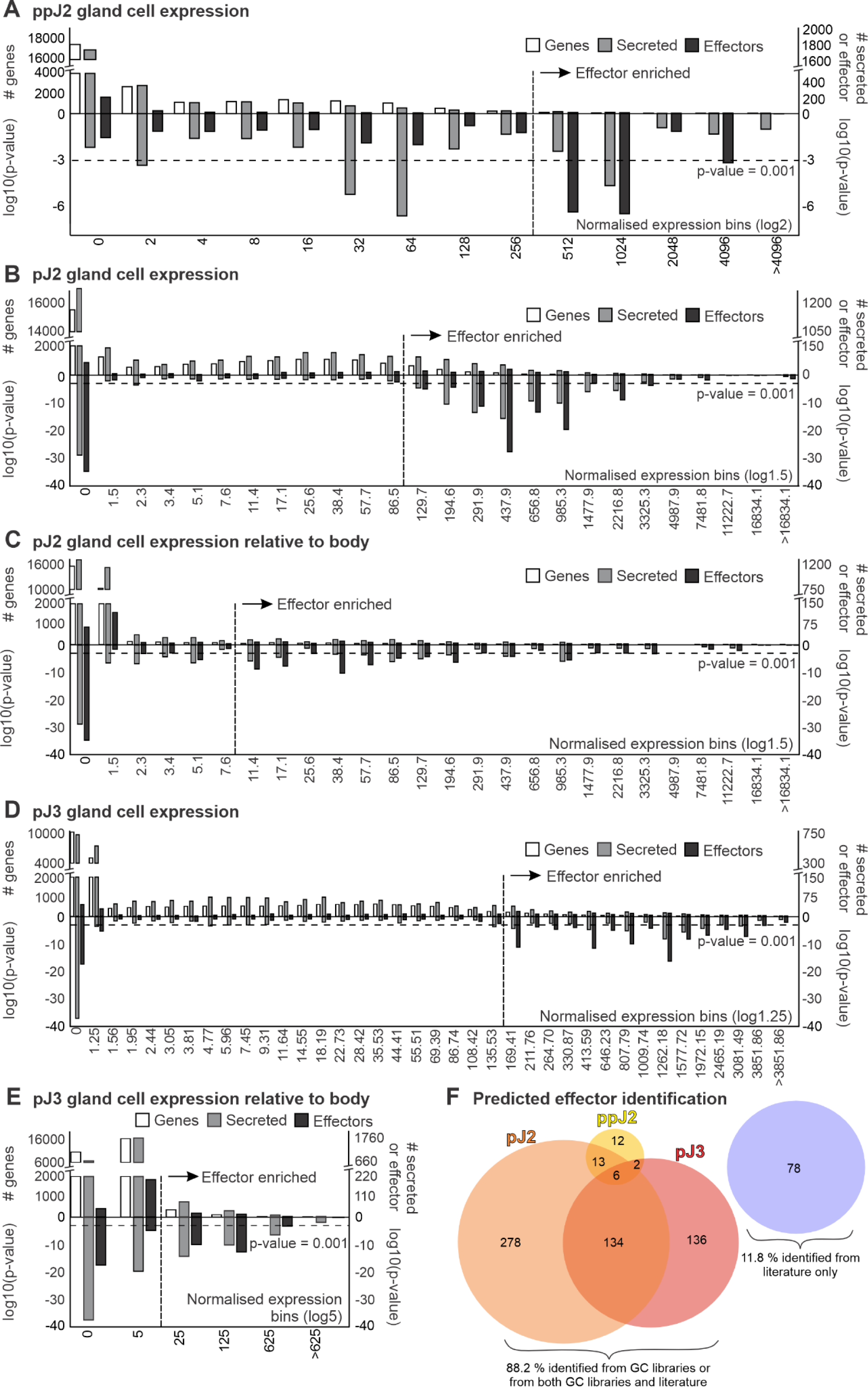
Gland cell enrichment of effectors and putatively secreted proteins. A-E) Effector enrichment in gland cell libraries. Putative effectors were identified using enrichment of effector-annotated genes and putative secreted proteins to identify an expression cutoff above which putatively secreted proteins are likely effectors. The upper axis shows the total number of genes (left) and effector-annotated genes and putative secreted proteins (right) in each expression bin (i.e. at each expression level). Hypergeometric distribution tests were used to determine either the enrichment or depletion of effectors in each bin. The lower axis shows the p-values from these tests. The horizontal dashed line denotes a p-value of 0.001. The vertical dashed line denotes the threshold expression level above which effector genes and or secreted proteins are largely or consistently enriched. **F**) Proportional Venn-diagram showing which gland cell libraries putative effectors were identified from.

**Figure S2.**
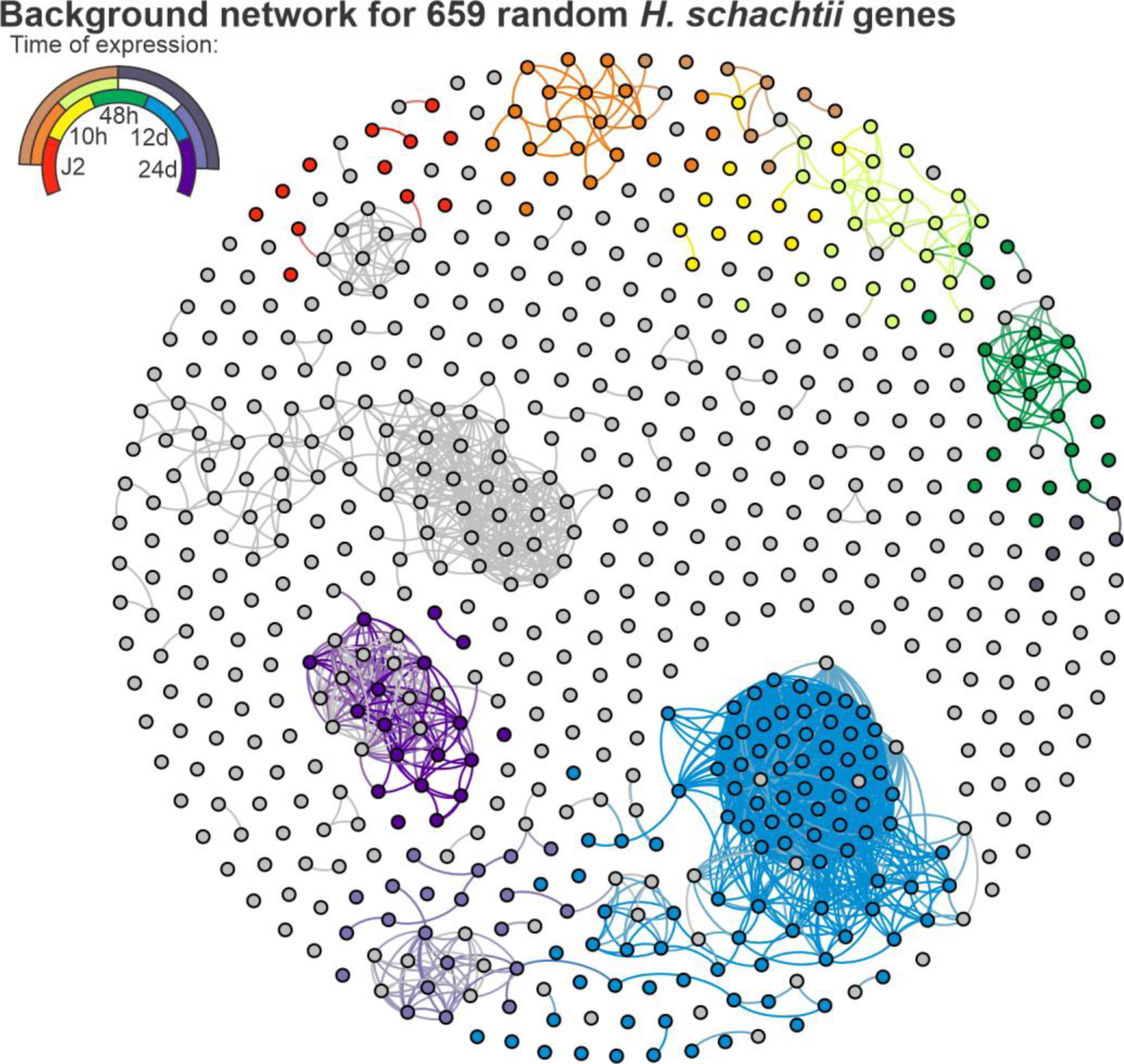
Transcriptional network of a random set of 659 genes. A transcriptional network of a random set of 659 *Heterodera schachtii* genes. Each circle represents one locus, and connections between circles indicate a correlation in expression of 0.975 or above (distance correlation coefficient) across the life cycle. The key indicates the expression supercluster as defined in (Siddique et al. 2022) - where, for example, genes with expression peaking at J2 are shown in red, 10 hours post infection in yellow, and J2_10 hours post infection shown in orange.

**Figure S3.**
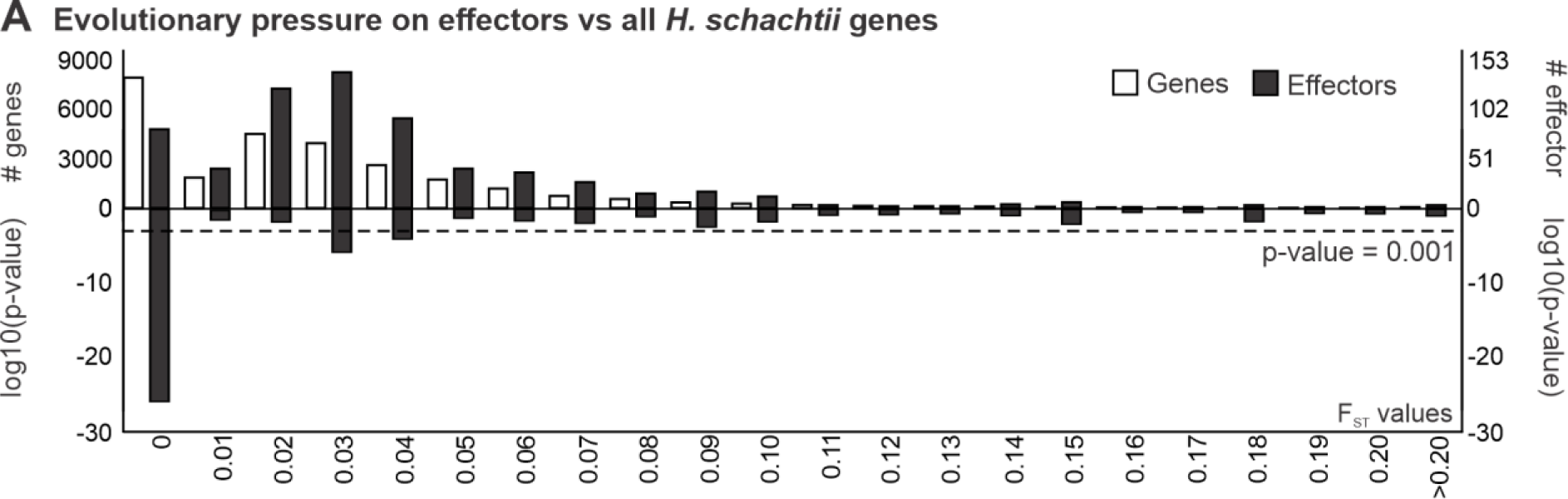
F_ST_ of effectors vs all genes. Frequency distribution of all *H. schachtii* genes (left axis) and effectors (right axis) across F_ST_ bins. Hypergeometric distribution tests were used to determine the enrichment or depletion of effectors in each bin. The lower axis shows the p-values from these tests. The horizontal dashed line denotes a p-value of 0.001.

**Figure S4.**
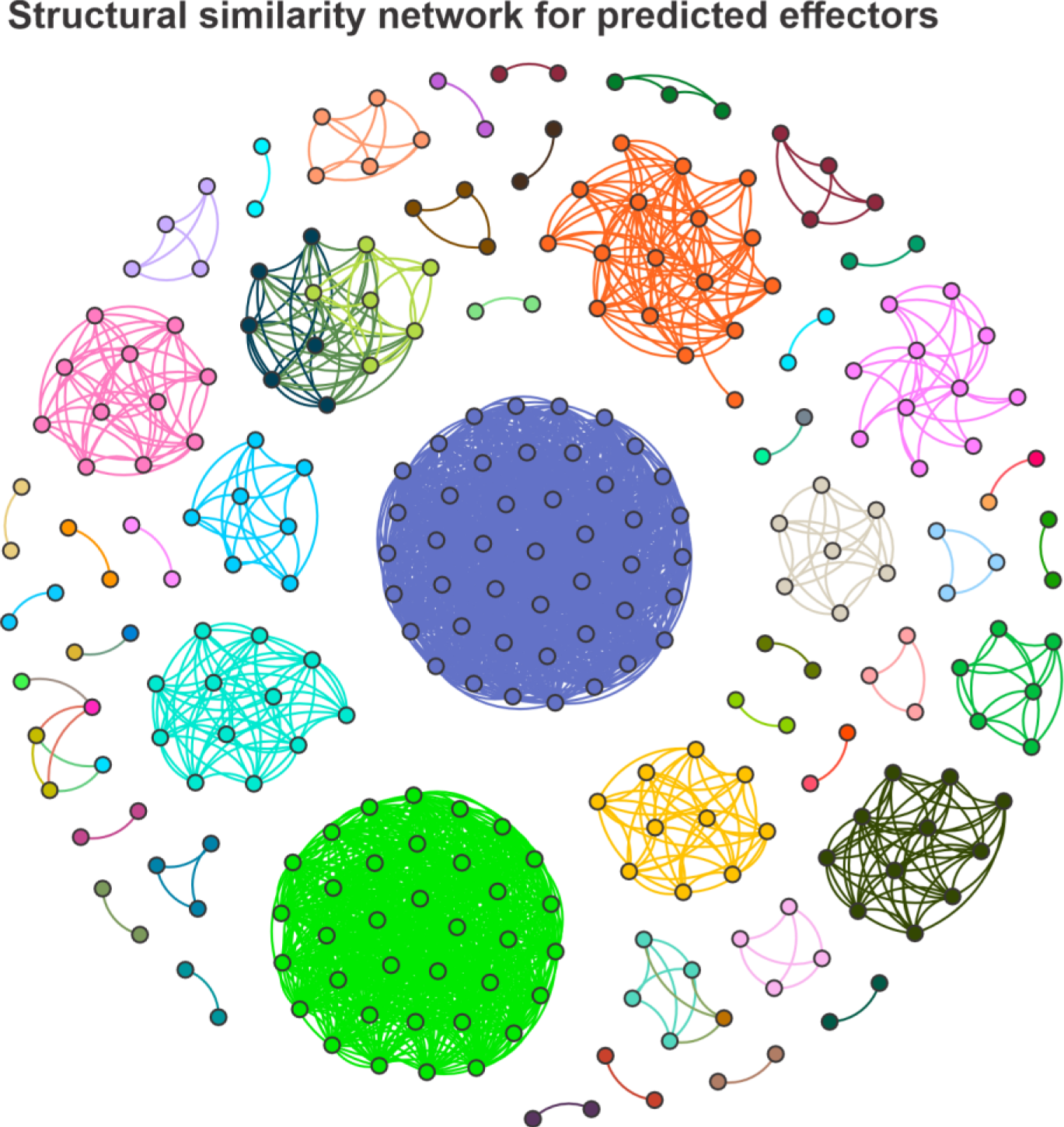
Structural similarity network for folded putative effectors. A structural similarity network for the predicted structures of putative *H. schachtii* effectors. Each circle represents one foldable effector gene locus (i.e. a fold with an average pLDDT > 50 and pTM > 0.5), and connections between circles indicate structural similarity TM-score >0.5 or above as determined using structure-based BLAST, Foldseek (Hutson 2023)). Colours indicate effector families as assigned in this study (Table S1).

**Figure S5.**
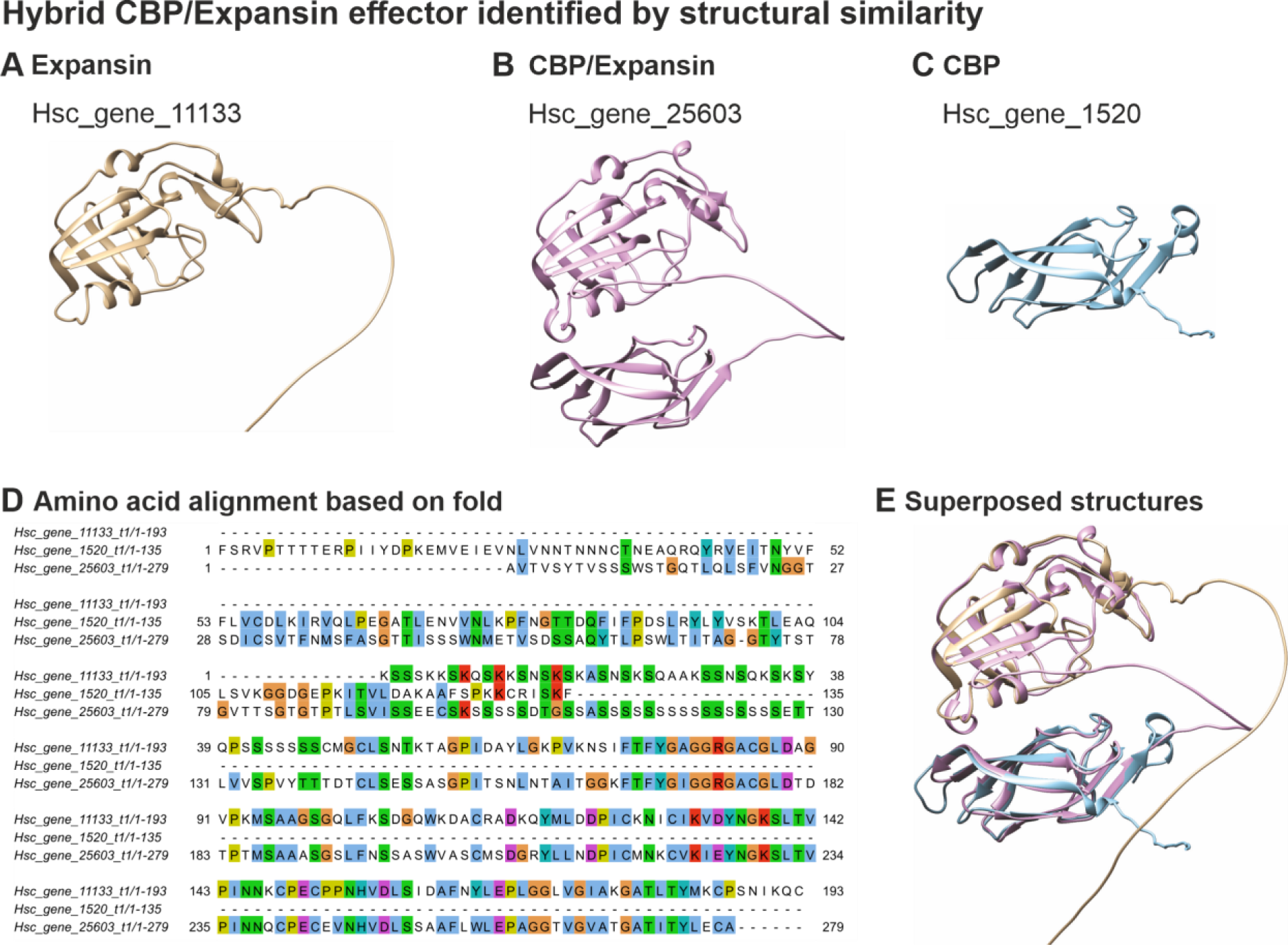
Hybrid CBP/Expansion effector identified by structural similarity of folded putative effectors. Predicted structures of *H. schachtii* effectors, showing **A)** an effector with an Expansin domain only, **B)** a hybrid effector with both a cellulose binding protein (CBP) domain and an Expansin domain, and **C)** an effector with a CBP domain only. Protein structures were predicted using ColabFold (using AlphaFold). **D)** An amino acid alignment of the three folded effectors. **E)** The superposed structures of expansin and CBP effectors onto hybrid CBP/Expansin effector.

**Figure S4.**
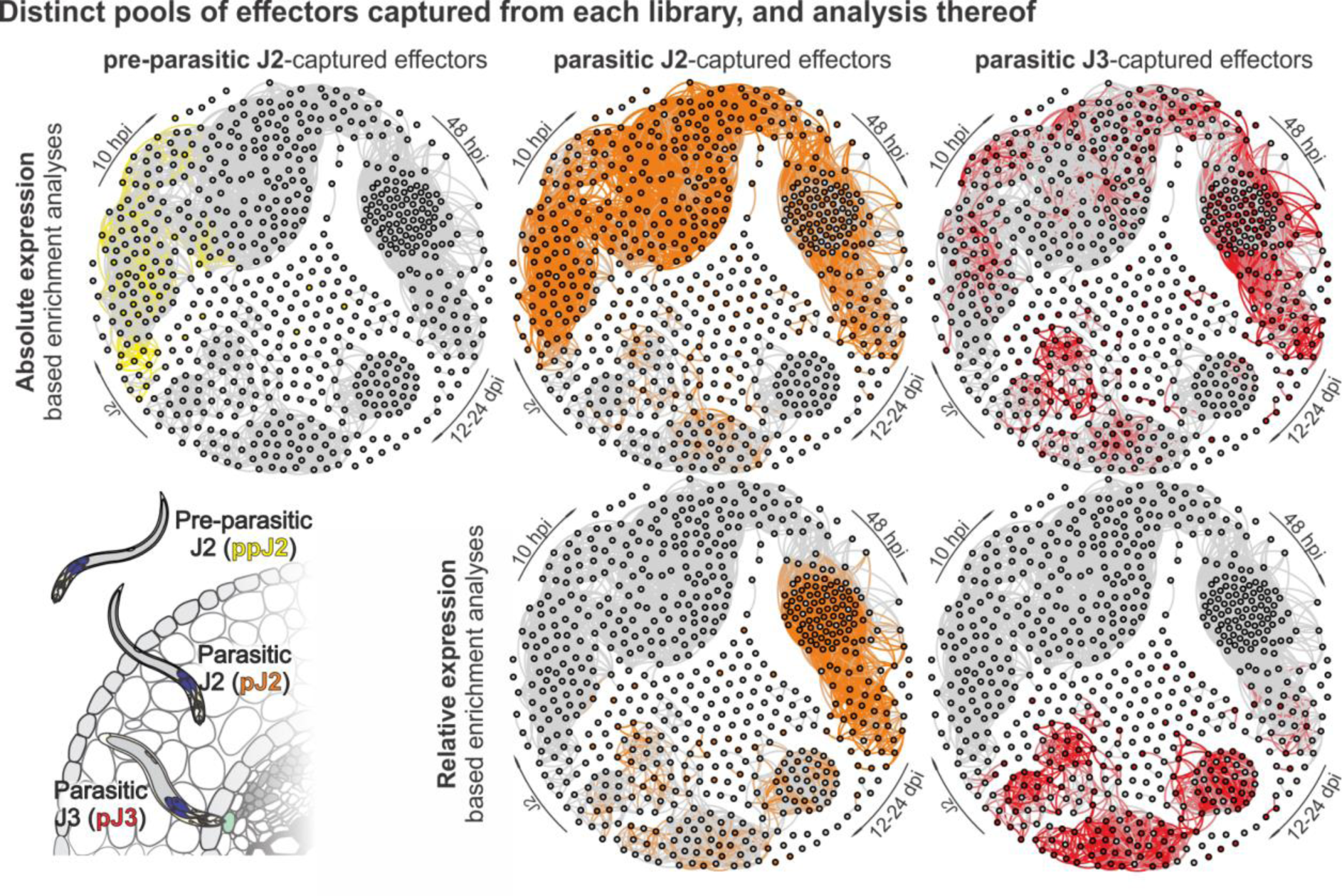
Effector enrichment analysis by library. Putative effectors were identified using enrichment of effector-annotated genes and putative secreted proteins to identify an expression cutoff above which putatively secreted proteins are likely effectors. For each of the pre-parasitic J2 (yellow), parasitic J2 (orange), and parasitic J3 (red) life-stages, the effectors that were identified are mapped to the network. For parasitic J2 and parasitic J3, both absolute expression-based analysis (top) and relative expression-based analysis (i.e. expression of each gene relativised to expression of said gene across the whole nematode body at the corresponding life-stage, bottom) are shown.

## References

Anders, Simon, Paul Theodor Pyl, and Wolfgang Huber. 2015. “HTSeq--a Python Framework to Work with High-Throughput Sequencing Data.” Bioinformatics 31 (2): 166–69.

“Babraham Bioinformatics - FastQC A Quality Control Tool for High Throughput Sequence Data.” n.d. Accessed May 26, 2023. https://www.bioinformatics.babraham.ac.uk/projects/fastqc/.

Barrio-Hernandez, Inigo, Jingi Yeo, Jürgen Jänes, Milot Mirdita, Cameron L. M. Gilchrist, Tanita Wein, Mihaly Varadi, Sameer Velankar, Pedro Beltrao, and Martin Steinegger. 2023. “Clustering Predicted Structures at the Scale of the Known Protein Universe.” Nature 622 (7983): 637–45.

Bastian, Mathieu, Sebastien Heymann, and Mathieu Jacomy. n.d. “Gephi: An Open Source Software for Exploring and Manipulating Networks.” Accessed December 11, 2023. https://gephi.org/publications/gephi-bastian-feb09.pdf.

Bbduk.Sh at Master · BioInfoTools/BBMap. n.d. Github. Accessed May 29, 2023. https://github.com/BioInfoTools/BBMap.

Bedell, Victoria M., Ying Wang, Jarryd M. Campbell, Tanya L. Poshusta, Colby G. Starker, Randall G. Krug 2nd, Wenfang Tan, et al. 2012. “In Vivo Genome Editing Using a High-Efficiency TALEN System.” Nature 491 (7422): 114–18.

Berardini, Tanya Z., Suparna Mundodi, Leonore Reiser, Eva Huala, Margarita Garcia-Hernandez, Peifen Zhang, Lukas A. Mueller, et al. 2004. “Functional Annotation of the Arabidopsis Genome Using Controlled Vocabularies.” Plant Physiology 135 (2): 745–55.

Bird, Alan F. 1983. “Changes in the Dimensions of the Oesophageal Glands in Root-Knot Nematodes during the Onset of Parasitism.” International Journal for Parasitology 13 (4): 343–48.

Boer, J. M. de, Y. Yan, G. Smant, E. L. Davis, and T. J. Baum. 1998. “In-Situ Hybridization to Messenger RNA in Heterodera Glycines.” Journal of Nematology 30 (3): 309–12.

Bolger, Anthony M., Marc Lohse, and Bjoern Usadel. 2014. “Trimmomatic: A Flexible Trimmer for Illumina Sequence Data.” *Bioinformatics*, btu170.

Cotton, James A., Catherine J. Lilley, Laura M. Jones, Taisei Kikuchi, Adam J. Reid, Peter Thorpe, Isheng J. Tsai, et al. 2014. “The Genome and Life-Stage Specific Transcriptomes of Globodera Pallida Elucidate Key Aspects of Plant Parasitism by a Cyst Nematode.” Genome Biology 15 (3): R43.

David S. Guttman, Boris A. Vinatzer, Sara F. Sarkar, Max V. Ranall, Gregory Kettler, and Jean T. Greenberg. 2002. “A Functional Screen for the Type III (Hrp) Secretome of the Plant Pathogen Pseudomonas Syringae.” Science 295 (5560): 1722–26.

Derevnina, Lida, Mauricio P. Contreras, Hiroaki Adachi, Jessica Upson, Angel Vergara Cruces, Rongrong Xie, Jan Skłenar, et al. 2021. “Plant Pathogens Convergently Evolved to Counteract Redundant Nodes of an NLR Immune Receptor Network.” PLoS Biology 19 (8): e3001136.

Dobin, Alexander, Carrie A. Davis, Felix Schlesinger, Jorg Drenkow, Chris Zaleski, Sonali Jha, Philippe Batut, Mark Chaisson, and Thomas R. Gingeras. 2013. “STAR: Ultrafast Universal RNA-Seq Aligner.” Bioinformatics 29 (1): 15–21.

Espada, Margarida, Sebastian Eves-van den Akker, Tom Maier, Paramasivan Vijayapalani, Thomas Baum, Manuel Mota, and John T. Jones. 2018. “STATAWAARS: A Promoter Motif Associated with Spatial Expression in the Major Effector-Producing Tissues of the Plant-Parasitic Nematode Bursaphelenchus Xylophilus.” BMC Genomics 19 (1): 553.

Eves-van den Akker, Sebastian, and Paul R. J. Birch. 2016. “Opening the Effector Protein Toolbox for Plant-Parasitic Cyst Nematode Interactions.” Molecular Plant 9 (11): 1451–53.

Eves-van den Akker, Sebastian, Dominik R. Laetsch, Peter Thorpe, Catherine J. Lilley, Etienne G. J. Danchin, Martine Da Rocha, Corinne Rancurel, Nancy E. Holroyd, James A. Cotton, and Amir Szitenberg. 2016. “The Genome of the Yellow Potato Cyst Nematode, Globodera Rostochiensis, Reveals Insights into the Basis of Parasitism and Virulence.” Genome Biology 17 (1): 124.

Eves-Van Den Akker, Sebastian, Catherine J. Lilley, Hazijah B. Yusup, John T. Jones, and Peter E. Urwin. 2016. “Functional C-TERMINALLY ENCODED PEPTIDE (CEP) Plant Hormone Domains Evolved de Novo in the Plant Parasite Rotylenchulus Reniformis.” Molecular Plant Pathology 17 (8): 1265–75.

Frei Dit Frey, Nicolas, and Bruno Favery. 2021. “Plant-Parasitic Nematode Secreted Peptides Hijack a Plant Secretory Pathway.” The New Phytologist.

Golinowski, W., F. M. W. Grundler, and M. Sobczak. 1996. “Changes in the Structure OfArabidopsis Thaliana during Female Development of the Plant-Parasitic NematodeHeterodera Schachtii.” Protoplasma 194 (1): 103–16.

Grundler, Florian M. W., Miroslaw Sobczak, and Wladyslaw Golinowski. 1998. “Formation of Wall Openings in Root Cells of Arabidopsis Thaliana Following Infection by the Plant-Parasitic Nematode Heterodera Schachtii.” European Journal of Plant Pathology / European Foundation for Plant Pathology 104 (6): 545–51.

Grynberg, Priscila, Roberto Coiti Togawa, Leticia Dias de Freitas, Jose Dijair Antonino, Corinne Rancurel, Marcos Mota do Carmo Costa, Maria Fatima Grossi-de-Sa, et al. 2020. “Comparative Genomics Reveals Novel Target Genes towards Specific Control of Plant-Parasitic Nematodes.” Genes 11 (11). 10.3390/genes11111347.

Guillen, Karine de, Diana Ortiz-Vallejo, Jérome Gracy, Elisabeth Fournier, Thomas Kroj, and André Padilla. 2015. “Structure Analysis Uncovers a Highly Diverse but Structurally Conserved Effector Family in Phytopathogenic Fungi.” PLoS Pathogens 11 (10): e1005228.

Guo, Xiaoli, Jianying Wang, Michael Gardner, Hiroo Fukuda, Yuki Kondo, J. Peter Etchells, Xiaohong Wang, and Melissa Goellner Mitchum. 2017. “Identification of Cyst Nematode B-Type CLE Peptides and Modulation of the Vascular Stem Cell Pathway for Feeding Cell Formation.” PLoS Pathogens 13 (2): e1006142.

Hogenhout, Saskia A., Renier A. L. Van der Hoorn, Ryohei Terauchi, and Sophien Kamoun. 2009. “Emerging Concepts in Effector Biology of Plant-Associated Organisms.” Molecular Plant-Microbe Interactions: MPMI 22 (2): 115–22.

Hu, Hui, Ya-Ru Miao, Long-Hao Jia, Qing-Yang Yu, Qiong Zhang, and An-Yuan Guo. 2019. “AnimalTFDB 3.0: A Comprehensive Resource for Annotation and Prediction of Animal Transcription Factors.” Nucleic Acids Research 47 (D1): D33–38.

Hussey, R. S., and C. W. Mims. 1990. “Ultrastructure of Esophageal Glands and Their Secretory Granules in the Root-Knot NematodeMeloidogyne Incognita.” Protoplasma 156 (1): 9–18.

Hutson, Matthew. 2023. “Foldseek Gives AlphaFold Protein Database a Rapid Search Tool.” *Nature*, June. 10.1038/d41586-023-02205-4.

Jiang, Rays H. Y., Sucheta Tripathy, Francine Govers, and Brett M. Tyler. 2008. “RXLR Effector Reservoir in Two *Phytophthora* Species Is Dominated by a Single Rapidly Evolving Superfamily with More than 700 Members.” Proceedings of the National Academy of Sciences of the United States of America 105 (12): 4874–79.

Jones, John T., Annelies Haegeman, Etienne G. J. Danchin, Hari S. Gaur, Johannes Helder, Michael G. K. Jones, Taisei Kikuchi, et al. 2013. “Top 10 Plant-Parasitic Nematodes in Molecular Plant Pathology.” Molecular Plant Pathology 14 (9): 946–61.

Jones, M. G. K. 1981. “Host Cell Responses to Endoparasitic Nematode Attack: Structure and Function of Giant Cells and Syncytia.” The Annals of Applied Biology 97 (3): 353–72.

Jumper, John, Richard Evans, Alexander Pritzel, Tim Green, Michael Figurnov, Olaf Ronneberger, Kathryn Tunyasuvunakool, et al. 2021. “Highly Accurate Protein Structure Prediction with AlphaFold.” Nature 596 (7873): 583–89.

Kempen, Michel van, Stephanie S. Kim, Charlotte Tumescheit, Milot Mirdita, Jeongjae Lee, Cameron L. M. Gilchrist, Johannes Söding, and Martin Steinegger. 2023. “Fast and Accurate Protein Structure Search with Foldseek.” *Nature Biotechnology*, May, 1–4.

Kummerfeld, Sarah K., and Sarah A. Teichmann. 2006. “DBD: A Transcription Factor Prediction Database.” Nucleic Acids Research 34 (Database issue): D74–81.

Lewis, S. E., S. M. J. Searle, N. Harris, M. Gibson, V. Lyer, J. Richter, C. Wiel, et al. 2002. “Apollo: A Sequence Annotation Editor.” Genome Biology 3 (12): RESEARCH0082.

Li, Gang, Nawaraj Dulal, Ziwen Gong, and Richard A. Wilson. 2023. “Unconventional Secretion of Magnaporthe Oryzae Effectors in Rice Cells Is Regulated by TRNA Modification and Codon Usage Control.” Nature Microbiology 8 (9): 1706–16.

Lilley, Catherine J., Abbas Maqbool, Duqing Wu, Hazijah B. Yusup, Laura M. Jones, Paul R. J. Birch, Mark J. Banfield, Peter E. Urwin, and Sebastian Eves-van den Akker. 2018. “Effector Gene Birth in Plant Parasitic Nematodes: Neofunctionalization of a Housekeeping Glutathione Synthetase Gene.” PLoS Genetics 14 (4): e1007310.

Lin, Zeming, Halil Akin, Roshan Rao, Brian Hie, Zhongkai Zhu, Wenting Lu, Nikita Smetanin, et al. 2023. “Evolutionary-Scale Prediction of Atomic-Level Protein Structure with a Language Model.” Science 379 (6637): 1123–30.

Love, Michael I., Wolfgang Huber, and Simon Anders. 2014. “Moderated Estimation of Fold Change and Dispersion for RNA-Seq Data with DESeq2.” Genome Biology 15 (12): 550.

Lovelace, Amelia H., Sara Dorhmi, Michelle T. Hulin, Yufei Li, John W. Mansfield, and Wenbo Ma. 2023. “Effector Identification in Plant Pathogens.” Phytopathology 113 (4): 637–50.

Maier, Tom R., Tarek Hewezi, Jiqing Peng, and Thomas J. Baum. 2013. “Isolation of Whole Esophageal Gland Cells from Plant-Parasitic Nematodes for Transcriptome Analyses and Effector Identification.” Molecular Plant-Microbe Interactions: MPMI 26 (1): 31– 35.

Maier, Tom R., Rick E. Masonbrink, Paramasivan Vijayapalani, Michael Gardner, Amanda D. Howland, Melissa G. Mitchum, and Thomas J. Baum. 2021. “Esophageal Gland RNA-Seq Resource of a Virulent and Avirulent Population of the Soybean Cyst Nematode.” Molecular Plant-Microbe Interactions: MPMI 34 (9): 1084–87.

Masonbrink, Rick, Tom R. Maier, Usha Muppirala, Arun S. Seetharam, Etienne Lord, Parijat S. Juvale, Jeremy Schmutz, et al. 2019. “The Genome of the Soybean Cyst Nematode (Heterodera Glycines) Reveals Complex Patterns of Duplications Involved in the Evolution of Parasitism Genes.” BMC Genomics 20 (1): 1–14.

Mirdita, Milot, Konstantin Schütze, Yoshitaka Moriwaki, Lim Heo, Sergey Ovchinnikov, and Martin Steinegger. 2022. “ColabFold: Making Protein Folding Accessible to All.” Nature Methods 19 (6): 679–82.

Molloy, Beth, Thomas Baum, and Sebastian Eves-van den Akker. 2023. “Unlocking the Development- and Physiology-Altering ‘effector Toolbox’ of Plant-Parasitic Nematodes.” Trends in Parasitology 39 (9): 732–38.

Nicol, J. M., S. J. Turner, D. L. Coyne, L. den Nijs, S. Hockland, and Z. Tahna Maafi. 2011. “Current Nematode Threats to World Agriculture.” In Genomics and Molecular Genetics of Plant-Nematode Interactions, edited by John Jones, Godelieve Gheysen, and Carmen Fenoll, 21–43. Dordrecht: Springer Netherlands.

Pearson, W. R. 2005. “Ranbpm Homologue Genes Characterised in the Cyst Nematodes Globodera Pallida and Globodera ‘Mexicana.’” Physiological and Molecular Plant Pathology 67 (1): 15–22.

Petersen, Thomas Nordahl, Soren Brunak, Gunnar von Heijne, and Henrik Nielsen. 2011. “SignalP 4.0: Discriminating Signal Peptides from Transmembrane Regions.” Nature Methods 8 (10): 785–86.

Pogorelko, Gennady, Jianying Wang, Parijat S. Juvale, Melissa G. Mitchum, and Thomas J. Baum. 2020. “Screening Soybean Cyst Nematode Effectors for Their Ability to Suppress Plant Immunity.” Molecular Plant Pathology 21 (9): 1240–47.

Robertson, L., W. M. Robertson, and J. T. Jones. 1999. “Direct Analysis of the Secretions of the Potato Cyst Nematode Globodera Rostochiensis.” Parasitology 119 (Pt 2) (August): 167–76.

Siddique, Shahid, Zoran S. Radakovic, Clarissa Hiltl, Clement Pellegrin, Thomas J. Baum, Helen Beasley, Andrew F. Bent, et al. 2022. “The Genome and Lifestage-Specific Transcriptomes of a Plant-Parasitic Nematode and Its Host Reveal Susceptibility Genes Involved in Trans-Kingdom Synthesis of Vitamin B5.” Nature Communications 13 (1): 6190.

Sperschneider, Jana, Donald M. Gardiner, Peter N. Dodds, Francesco Tini, Lorenzo Covarelli, Karam B. Singh, John M. Manners, and Jennifer M. Taylor. 2016. “EffectorP: Predicting Fungal Effector Proteins from Secretomes Using Machine Learning.” The New Phytologist 210 (2): 743–61.

Steenbrugge, Joris J. M. van, Sven van den Elsen, Martijn Holterman, Jose L. Lozano-Torres, Vera Putker, Peter Thorpe, Aska Goverse, Mark G. Sterken, Geert Smant, and Johannes Helder. 2023. “Comparative Genomics among Cyst Nematodes Reveals Distinct Evolutionary Histories among Effector Families and an Irregular Distribution of Effector-Associated Promoter Motifs.” Molecular Ecology 32 (6): 1515– 29.

Steinegger, Martin, and Johannes Söding. 2017. “MMseqs2 Enables Sensitive Protein Sequence Searching for the Analysis of Massive Data Sets.” Nature Biotechnology 35 (11): 1026–28.

Teufel, Felix, José Juan Almagro Armenteros, Alexander Rosenberg Johansen, Magnús Halldór Gíslason, Silas Irby Pihl, Konstantinos D. Tsirigos, Ole Winther, Søren Brunak, Gunnar von Heijne, and Henrik Nielsen. 2022. “SignalP 6.0 Predicts All Five Types of Signal Peptides Using Protein Language Models.” Nature Biotechnology 40 (7): 1023.

Vieira, Paulo, Thomas R. Maier, Sebastian Eves-van den Akker, Dana K. Howe, Inga Zasada, Thomas J. Baum, Jonathan D. Eisenback, and Kathryn Kamo. 2018. “Identification of Candidate Effector Genes of Pratylenchus Penetrans.” *Molecular Plant Pathology*, February. 10.1111/mpp.12666.

Vieira, Paulo, Roxana Y. Myers, Clement Pellegrin, Catherine Wram, Cedar Hesse, Thomas R. Maier, Jonathan Shao, et al. 2021. “Targeted Transcriptomics Reveals Signatures of Large-Scale Independent Origins and Concerted Regulation of Effector Genes in Radopholus Similis.” PLoS Pathogens 17 (11): e1010036.

Vieira, Paulo, Jonathan Shao, Paramasivan Vijayapalani, Thomas R. Maier, Clement Pellegrin, Sebastian Eves-van den Akker, Thomas J. Baum, and Lev G. Nemchinov. 2020. “A New Esophageal Gland Transcriptome Reveals Signatures of Large Scale de Novo Effector Birth in the Root Lesion Nematode Pratylenchus Penetrans.” BMC Genomics 21 (1): 738.

Wickham, Hadley, Mara Averick, Jennifer Bryan, Winston Chang, Lucy McGowan, Romain François, Garrett Grolemund, et al. 2019. “Welcome to the Tidyverse.” Journal of Open Source Software 4 (43): 1686.

